# Molecular basis for chikungunya virus recognition of a mosquito-specific receptor

**DOI:** 10.64898/2026.07.28.741352

**Authors:** Xiaoyi Fan, Wanyu Li, Jesse S. Plung, Jessica A. Plante, Ellie M. Hajovsky, Nisha Tapryal, Jennifer Diaz, Nicholas C. Hazell, Chenggong Ji, Zhiqi Liu, Catherine E. Hammond, Scott C. Weaver, Kenneth S. Plante, Jonathan Abraham

## Abstract

Alphaviruses are arthropod-borne viruses that recognize cellular receptors in both vertebrate hosts and mosquito vectors to complete their transmission cycle, yet how they maintain recognition of receptors across evolutionarily divergent host species remains unresolved. Among alphaviruses, chikungunya virus (CHIKV), which is primarily vectored in urban settings by *Aedes* species mosquitoes, is the most widespread, and causes explosive outbreaks that can involve hundreds of thousands to millions of cases annually. The cell adhesion protein Lachesin is a mosquito-specific cellular receptor for CHIKV and multiple other arthritogenic alphaviruses. The envelope E2–E1 glycoproteins of these alphaviruses broadly recognize Lachesin orthologs from diverse mosquito species, but not other insects or arachnids. Lachesin genetic manipulation to prevent mosquito virus infection without interfering with endogenous receptor function could have a major impact for CHIKV control. Here, we determined high-resolution cryo-electron microscopy (cryo-EM) structures of alphaviruses bound to *Aedes albopictus* Lachesin. Comparative analysis of Lachesin-bound CHIKV, Semliki Forest virus (SFV), and Middelburg virus (MIDV) revealed that these three genetically divergent viruses use a similar surface to recognize Lachesin domain 1, but with reorganized E2–E1 glycoprotein contact residues. We show that a soluble *Ae. albopictus* Lachesin receptor decoy protein blocks the E2–E1-mediated entry of CHIKV and other arthritogenic alphaviruses into mammalian cells with greater breadth than a vertebrate receptor MXRA8 decoy and protects against lethal SFV challenge and CHIKV pathogenesis in murine models. Additionally, we identified a naturally occurring single residue Lachesin polymorphism that is found in some *Anopheles* (malaria vector) mosquitoes, and fully ablates CHIKV E2–E1 recognition, informing strategies for mosquito-targeted genetic interventions that could prevent mosquito vector infection and virus transmission. These findings define distinct determinants of receptor binding in mosquitoes and humans for arthritogenic alphaviruses, with implications for countermeasure development and outbreak preparedness.

## Introduction

Arthritogenic alphaviruses are mosquito-borne pathogens that cause febrile illnesses accompanied by rash, arthralgia, and arthritis in humans (*6*). They include chikungunya virus (CHIKV), o’nyong-nyong virus (ONNV), Mayaro virus (MAYV), and Ross River virus (RRV), which are all members of the Semliki Forest (SF) complex, and Barmah Forest virus (BFV) of the Barmah Forest complex (Figure 1A) (*6*). Among them, CHIKV is the most widespread globally and infects an estimated 16.9 million annually, while 2.8 billion are at risk of infection in 119 countries (*7-9*).

**Figure 1.**
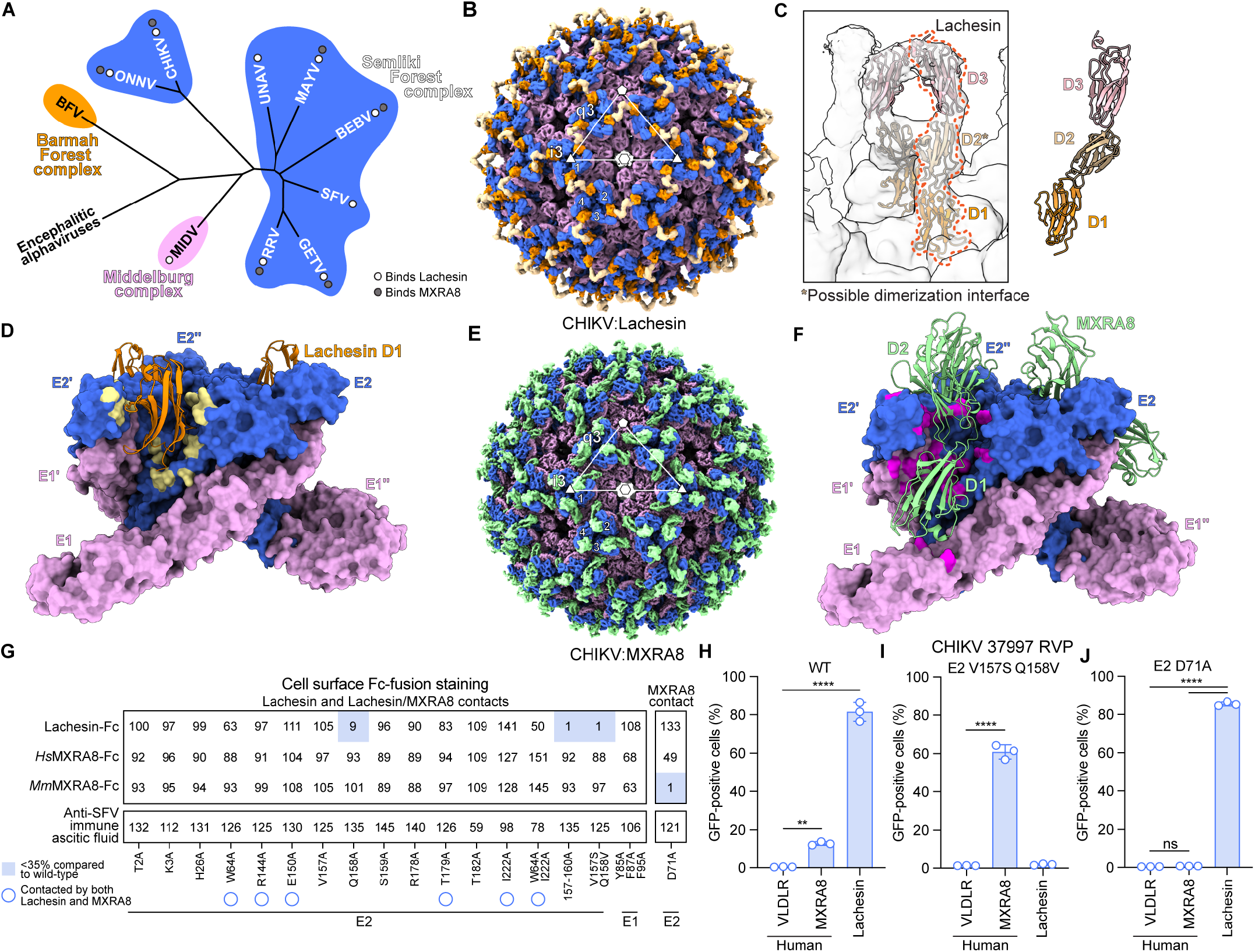
Molecular determinants of CHIKV interactions with mosquito and mammalian receptors. **(A)** Phylogenetic tree of the indicated alphaviruses based on sequence alignment of their E2 glycoproteins. Encephalitic alphaviruses: WEEV, VEEV and EEEV were used as an outgroup (not shown). Viruses that can use mosquito Lachesin or the mammalian receptor MXRA8 are denoted (*1, 2*). **(B)** Cryo-EM map of Lachesin–Fc bound to CHIKV VLP. The E2 and E1 glycoproteins are shown in blue and pink, respectively, and the bound Lachesin D1 is in orange, with additional density colored beige. Icosahedral symmetry axes (i5, i3, i2) are indicated with a pentagon, triangle, and hexagon, respectively. The icosahedral threefold (i3) and quasi-threefold (q3) E2–E1 trimers are indicated. Four E2–E1 heterodimers that constitute an asymmetric unit are numbered. **(C)** Fitting of the Lachesin ectodomain structure predicted by AlphaFold3 (*3*) into the cryo-EM density map of CHIKV VLPs bound to Lachesin–Fc, contoured at a lower threshold reveals that Lachesin D1 is inserted into clefts formed by adjacent E2–E1 heterodimers. Asterisk indicates possible dimerization at the level of Lachesin D2. **(D)** CHIKV E2–E1 trimer bound to Lachesin D1. The E2–E1 trimer is surface-rendered and the bound Lachesin D1 molecules are shown as ribbon diagrams. CHIKV E2–E1 residues that interact with D1 are shown as a yellow surface. See Figure S3 for detailed viral glycoprotein–Lachesin interactions. **(E)** Cryo-EM map of *Hs*MXRA8–Fc bound to CHIKV VLP. Density for E2 and E1 is shown in blue and pink, respectively, and density for *Hs*MXRA8 is colored green. Icosahedral symmetry axes, the i3 and q3 E2–E1 trimers, and the four E2–E1 heterodimers that constitute an asymmetric unit are indicated as in **C. (F)** CHIKV E2–E1 trimer bound to *Hs*MXRA8. The CHIKV E2–E1 trimer is surface rendered, and bound *Hs*MXRA8 molecules are shown as ribbon diagrams. CHIKV E2–E1 residues that interact with *Hs*MXRA8 are shown in magenta. **(G)** Staining of HEK293T cells transfected to express wild-type or mutant CHIKV E3–E2–(6K/TF)–E1 proteins by the indicated Fc fusion proteins. See Methods and Figure S9A for additional information. **(H, I)** K562 cells ectopically expressing human VLDLR, human MXRA8, or *Ae. albopictus* Lachesin were infected with GFP-expressing RVPs bearing wild-type (WT) CHIKV 37997 E2–E1 at an MOI of 1 (measured on K562 cells expressing *Ae. aegypti* Lachesin) (**H**) or CHIKV 37997 E2–E1 containing the V157S Q158V E2 mutations at an MOI of 1 (measured on K562 cells expressing *Hs*MXRA8) (**I**). Infection was measured using flow cytometry. **(J)** K562 cells ectopically expressing human VLDLR, human MXRA8, or *Ae. albopictus* Lachesin were infected with CHIKV 37997 RVPs with E2–E1 containing the E2 D71A mutation at an MOI of 2 (measured on K562 cells expressing *Ae. albopictus* Lachesin). Infection was measured using flow cytometry. Data are mean from three experiments (*n* = 3) (**G**) or mean ± s.d. from three experiments performed in duplicates (*n* = 3) (**H, I, J**). One-way ANOVA with Dunnett’s multiple comparisons test, \*\**P* = 0.0035 (**D**) \*\*\*\**P* < 0.0001 (**H, I, J**).

First isolated in Tanzania in 1952, CHIKV has, on multiple occasions, expanded from its ancestral sylvatic transmission cycles in Africa into human-amplified global epidemics (*10-13*). In sub-Saharan Africa, CHIKV circulates in enzootic, sylvatic cycles, sequentially infecting non-human primates (and possibly other vertebrate hosts) and forest-dwelling *Aedes* mosquitoes (*11, 14-16*). CHIKV can emerge from its sylvatic reservoir when an infected enzootic or bridge vector bites a human host. In such cases, CHIKV is then transmitted among humans in urban environments by the anthropophilic and peridomestic *Ae. aegypti* and *Ae. albopictus* mosquitoes (*6, 7*). In recent years, CHIKV has expanded into previously unaffected regions, driven in part by globalization and climate-associated shifts in mosquito vector distribution, as well as vector-adaptive evolution (*17-19*). Although historically confined to the Old World, CHIKV reached the Americas in the 2010s, and in 2025, the virus experienced a marked global resurgence, with large-scale outbreaks on Indian Ocean islands, continued high transmission in the Americas and South Asia, and locally acquired cases reported in several temperate regions, including China, France, Italy, and the United States, including the first locally acquired case in New York State (*8, 20*).

In addition to CHIKV, the related SF complex alpha-viruses—ONNV, MAYV, and RRV—cause locally restricted outbreaks in Africa, the Americas, and Australia, respectively (*7*). However, these alphaviruses have the potential to emerge into new regions. In their enzootic cycles, these viruses are each transmitted to mammalian amplification hosts by infected mosquitoes of distinct genera, although the enzootic cycle of ONNV remains enigmatic.

The alphavirus genome encodes four nonstructural proteins (nsP1–nsP4) and six structural proteins (capsid, E3, E2, 6K, TF, and E1) (*21*). Virions have *T* = 4 icosahedral symmetry, with 240 E2–E1 heterodimers arranged into 80 trimeric glycoprotein spikes (*22-24*). E2–E1 glycoprotein heterodimers mediate receptor binding and membrane fusion that is triggered upon exposure to low pH in acidified endosomes.

The immunoglobulin superfamily cell adhesion protein matrix remodeling-associated 8 (MXRA8) is a receptor on mammalian cells for CHIKV, ONNV, MAYV, RRV and the veterinary pathogen Getah virus (GETV) (*2, 25*). The SF complex also contains Semliki Forest virus (SFV), which can bind very low-density lipoprotein receptor (VLDLR) and apolipoprotein E receptor 2 (ApoER2, also known as LRP8) as cellular receptors (*26-28*). Middelburg virus (MIDV), which is in the antigenically distinct Middelburg (MID) serocomplex and circulates enzootically in Africa causing fever and encephalitis in livestock, is closely related to SF complex alphaviruses (Figure 1A).

Although invertebrates lack an MXRA8 ortholog, we recently identified Lachesin, a glycosylphosphatidylinositol (GPI)-anchored immunoglobulin superfamily protein that is expressed on mosquito cells, as a receptor for multiple alphaviruses including CHIKV, SFV, ONNV, MAYV, RRV, GETV, and MIDV (*1*).

A lack of information on molecular determinants of arbovirus vector competency has prevented efforts to specifically target pathways in mosquitoes that would result in selective resistance to infection by specific arboviruses. Lachesin is expressed in the mosquito midgut, which is the first site of infection during an infectious bloodmeal. Obtaining a molecular understanding of how arthritogenic alphaviruses interact with Lachesin would afford the opportunity to selectively engineer mosquito resistance through changes in Lachesin that block virus receptor-mediated entry while otherwise preserving endogenous receptor function.

Here, we determined cryo-EM structures of multiple alphaviruses bound to Lachesin, clarifying how some alphaviruses have evolved the ability to recognize divergent cell surface receptors in mammalian hosts and mosquito vectors. We demonstrate that a Lachesin receptor decoy protein has broad-spectrum neutralizing activity in vitro that translates into protection in animal models of alphavirus pathogenesis. We identify a single residue polymorphism found in the Lachesin ortholog of *Anopheles funestus*, a malaria vector but not a CHIKV vector, that when introduced into *Ae. albopictus* Lachesin prevents CHIKV E2–E1-mediated infection. Our findings provide a molecular foundation for strategies targeting the *Lac* gene to render mosquito populations resistant to arthritogenic alphavirus infection.

## Results

### Structural basis for CHIKV recognition of Lachesin

Lachesin contains three immunoglobulin-like domains (D1–D3) in its ectodomain. We generated a human IgG1 Fc fusion protein containing *Ae. albopictus* Lachesin D1–D3 (Lachesin–Fc) (Figure S1A and S2A). In biolayer interferometry experiments, biosensors coated with CHIKV strain 37997 virus-like particles (VLPs) bound Lachesin–Fc in solution with high affinity (K_D_ of 5 nM) (Figs. S1B and S1C). We determined a cryo-EM structure of CHIKV VLPs in complex with *Ae. albopictus* Lachesin–Fc to 6.4 Å global resolution, with icosahedral symmetry (Figs. 1B, S1D, and Table S1). We fitted an AlphaFold 3.0 (AF3) (*29*)-predicted model of Lachesin D1–D3 into the low-contour density, which revealed D1 inserted into the cleft formed by two adjacent E2– E1 protomers (Figs. 1C, 1D, and S1E). We observed additional density, attributable to D2 and D3 extending away from the VLP, that did not contact E2–E1 glycoproteins but may participate in receptor homodimerization at the level of D2 (Fig. 1C). Prior domain mapping experiments had implicated Lachesin D1 as the binding site for CHIKV (*1*), consistent with this cryo-EM structure. Receptor homodimerization through D2 may indirectly enhance the strength of D1 interactions with E2–E1 and explain why overexpression in non-permissive cells of a construct that contains D1 and D2 more robustly supports CHIKV entry than does a construct that only contains D1 (*1*).

We used a block-based refinement strategy (*30*) to obtain a 3.4 Å map for the CHIKV:Lachesin–Fc complex asymmetric unit (Figs. S1D, S1F, and S1G). Lachesin D1 has an immunoglobulin-like β-sandwich fold comprising ten antiparallel β-strands (A–J) organized into two β-sheets and stabilized by a conserved disulfide bond between strands C and I (Fig. S1H). Lachesin D1 buries a surface area of ∼1500 Å^2^ in each cleft of the CHIKV E2–E1 trimer, comparable in size to the buried surface area observed between western equine encephalitis virus (WEEV) E2–E1 and the extracellular cadherin repeat 1 (EC1) of one of its receptors, protocadherin 10 (PCDH10) (∼1500 Å^2^) (Table S2) (*31*). One face of Lachesin contacts domain A and the β-ribbon connector of CHIKV E2, while the other face contacts domains A and B of the neighboring E2 protomer (E2′) (Figs. 1D and S3).

### Determinants of CHIKV mammalian vs. mosquito receptor binding

Previously reported structures of CHIKV bound to MXRA8 include a 4.1 Å cryo-EM structure of murine MXRA8 bound to CHIKV (*25*) and a 3.5 Å X-ray crystal structure of trimeric CHIKV E2–E1 bound to human MXRA8 (*32*). To allow for detailed comparison of molecular contacts that CHIKV makes with Lachesin and MXRA8, we determined a higher resolution cryo-EM structure of CHIKV VLPs bound to human MXRA8 (3.0 Å resolution of the receptor-bound VLP asymmetric unit) (Figs. 1E, 1F, and S4A–S4D). The CHIKV E2–E1 surface buried by one MXRA8 molecule is ∼2,200 Å^2^, which is much larger than the surface buried by one Lachesin molecule (∼1,500 Å^2^) (Table S2).

Some of the MXRA8 and Lachesin CHIKV E2 contact residues overlap (Fig. S5). To test the selective contribution of contact residues to Lachesin and MXRA8 binding, we next transfected HEK293T cells to express WT or mutant CHIKV E2–E1 glycoproteins containing alanine substitutions of Lachesin-contacting residues and stained cells with Lachesin–Fc, human MXRA8–Fc (*Hs*MXRA8–Fc), or murine MXRA8–Fc (*Mm*MXRA8–Fc) (Figs. 1G, S2B, S2C, S6A, S6B, and S9A). Lachesin–Fc bound robustly to cells transfected with WT CHIKV E2–E1, and most alanine substitutions of individual CHIKV E2 Lachesin contact residues minimally affected Lachesin–Fc staining (Fig. 1G), suggesting that the large CHIKV-Lachesin binding interface can tolerate disruption of single contacts.

However, alanine substitution of E2 residue Q158 resulted in a profound reduction in Lachesin–Fc staining (Fig. 1G). This E2 residue is in the central arch of the β-ribbon connector and is involved in a network of interactions that also includes E2 V157, E2 S159, and multiple Lachesin residues (Fig. S3C). The Q158 side chain does not interact with MXRA8; accordingly, the Q158A substitution did not impact *Hs*MXRA8– or *Mm*MXRA8–Fc staining (Fig. 1G). Simultaneous substitution of E2 residues V157–T160 with alanine completely abolished Lachesin–Fc staining with no impact on MXRA8–Fc staining. These results suggest that the central arch of the CHIKV E2 β-ribbon connector is critical for Lachesin engagement. By contrast, alanine substitution of CHIKV E2 residues whose side chains contact both Lachesin and MXRA8 had no effects on cell-surface staining by *Hs*MXRA8–Fc or *Mm*MXRA8–Fc.

A recent deep-mutational scanning (DMS) study reported that the E2 V157S and Q158V substitutions impair CHIKV entry into *Ae. albopictus* C6/36 cells but not HEK293T cells ectopically expressing human MXRA8 (*33*). We surmised that the E2 V157S and Q158V mutations selectively impact Lachesin binding by disrupting key contacts the CHIKV E2 β-ribbon connector makes with Lachesin (Fig. S3C), but not with MXRA8. Indeed, the E2 V157S/Q158V substitutions abolished Lachesin–Fc staining of HEK293T cells expressing this CHIKV E2–E1 glycoprotein mutant but had no effect on *Hs*MXRA8–Fc or *Mm*MXRA8–Fc staining (Fig. 1G).

To test the impact of the E2 V157S and Q158V substitutions on CHIKV E2–E1-mediated cellular entry, we also used single-cycle reporter virus particles (RVPs), which contain the RRV genome with the E3–E2–(6K/TF)–E1 coding sequence replaced with green fluorescent protein (GFP), and heterologous alphavirus glycoproteins on their surface provided in *trans* (*27*). RVPs bearing WT CHIKV E2– E1 (strain 37997) glycoproteins did not infect K562 cells expressing VLDLR (negative control), but infected cells expressing human MXRA8 and *Ae. albopictus* Lachesin (Figs. 1H, S6C, and S7A–C). Consistent with a selective inability to bind Lachesin, CHIKV 37997 RVPs harboring the E2 V157S and Q158V mutations infected K562 cells expressing human MXRA8, but not Lachesin (Fig. 1I).

We also sought to identify CHIKV E2–E1 determinants of MXRA8, but not Lachesin binding. Lachesin only engages CHIKV E2, whereas MXRA8 also contacts the E1’ fusion loop (Figs. 1F, S4E, and S4F). Alanine substitutions of E1’ fusion loop residues (Y85, F87, and F95) had little to no effect on MXRA8–Fc staining, and, as expected, no effect on Lachesin–Fc staining (Fig. 1G). The E1 glycoprotein fusion loop contacts are therefore not critically required for MXRA8 engagement. A prior study using cell-surface-displayed CHIKV E2–E1 reported that the D71A substitution in CHIKV E2 domain A ablates recognition by murine MXRA8–Fc (*2*); D71 was also identified as important for entry of CHIKV RVPs into human cells expressing MXRA8 but not for entry into C6/36 (*Ae. albopictus*) cells in a DMS study (*33*). E2 residue D71 is central to an extensive network of polar contacts with MXRA8 but does not contact Lachesin (Fig. S8). Binding of murine MXRA8–Fc, but not of Lachesin–Fc, was impaired on cells expressing CHIKV E2–E1 D71A (Fig. 1G). Binding of human MXRA8–Fc was only slightly impaired in our immunostaining experiments with cells expressing CHIKV E2–E1 D71A. However, CHIKV RVPs containing the D71A E2 substitution were unable to infect K562 cells expressing human MXRA8 but could still infect K562 cells expressing Lachesin (Fig. 1J).

Thus, while Lachesin and MXRA8 bind some overlapping residues on the CHIKV E2 glycoprotein, E2 contact residues that most critically contribute to affinity for Lachesin are largely separate from those that are critical for interactions with MXRA8.

### SFV receptor-binding sites are functionally separated

The E2 glycoproteins of CHIKV and SFV only share 58% amino acid sequence identity (Fig. 2A). To examine the basis for broad alphavirus recognition of Lachesin, we next determined a cryo-EM structure of *Ae. albopictus* Lachesin bound to SFV VLPs. We obtained a 4.9 Å map of an SFV VLP bound to Lachesin, and a 3.2 Å map of the asymmetric unit (Figs. 2B and S10). Although E3 is retained in SFV VLPs, the presence of E3 did not influence the Lachesin binding mode, and SFV engages Lachesin in a similar general binding mode as CHIKV (Figs. 2B–D). The individual virus glycoprotein– receptor contacts, however, are substantially reorganized when the CHIKV and SFV receptor-bound structures are compared (Figs. S3 and S11). Structural plasticity in individual receptor contacts thus supports a similar Lachesin-binding mode despite extensive sequence variation in the CHIKV and SFV spike glycoproteins.

**Figure 2.**
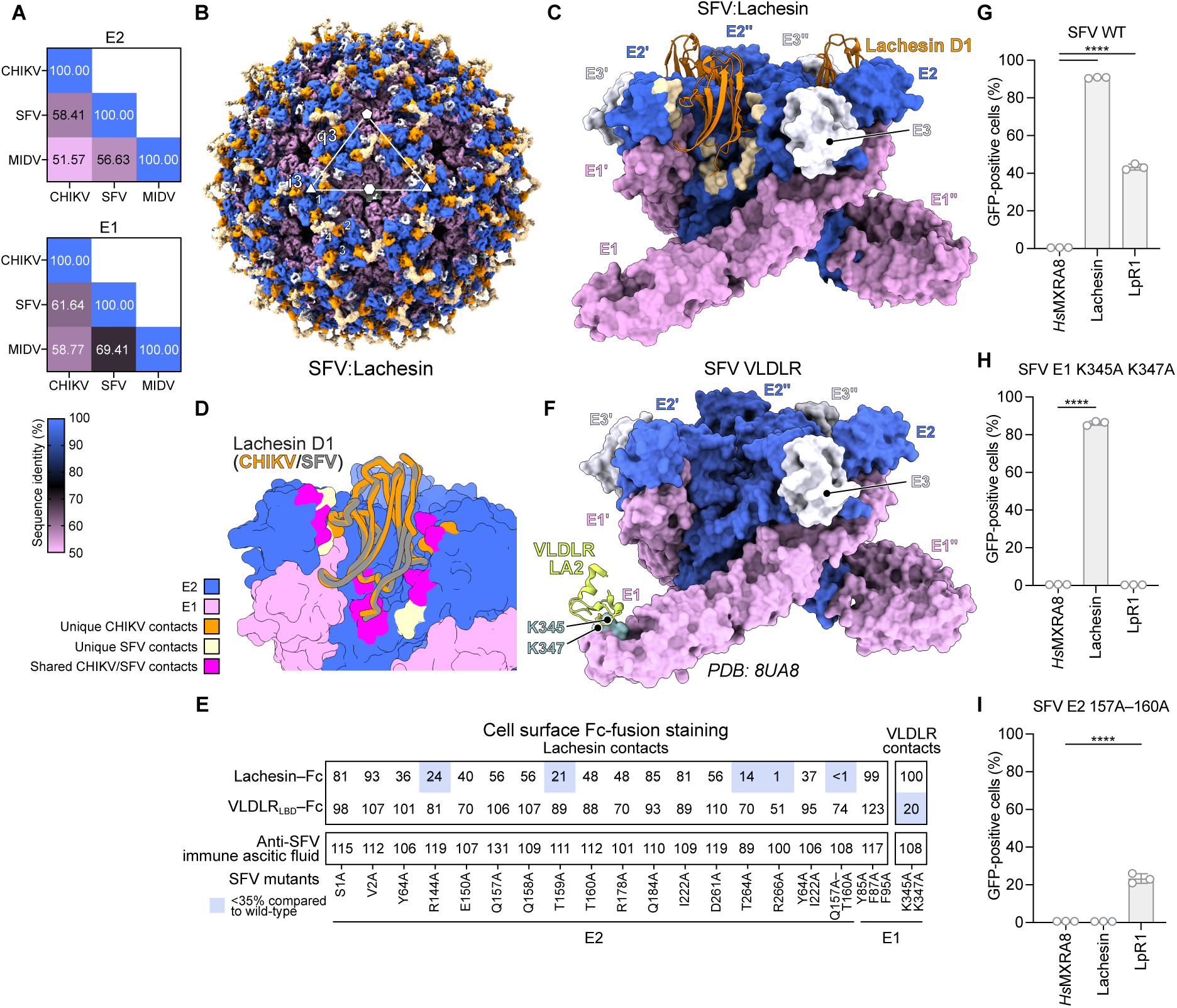
Molecular determinants of SFV interactions with Lachesin and VLDLR. **(A)** Amino acid identity matrices of CHIKV 37997, SFV SFV4, and MIDV MIDV857 E2 (top) or E1 (bottom) glycoproteins. **(B)** Overall cryo-EM map of Lachesin–Fc bound to SFV VLP. Density for the SFV E2 and E1 glycoproteins are shown in blue and pink, respectively. E3 density is shown in white. Density for bound Lachesin D1 is colored orange, while additional density is in beige. Icosahedral symmetry axes (i5, i3, i2) are indicated with a pentagon, triangle, and hexagon, respectively. The icosahedral i3 and q3 E2–E1 trimers are indicated. Four E2–E1 heterodimers that constitute an asymmetric unit are numbered. **(C)** SFV E2–E1 trimer bound to Lachesin D1. The SFV E2–E1 trimers and their associated E3 proteins are surface rendered, and Lachesin D1 molecules are shown as ribbon diagrams. SFV E2–E1 residues that interact with D1 are in beige. **(D)** Superposition of CHIKV E2–E1 and SFV E2–E1 bound to Lachesin D1, with structural superposition based on E2–E1 coordinates. E2 and E1 are shown in blue and pink, respectively. Shared contact residues on both CHIKV and SFV E2 are shown in magenta, contacts unique to CHIKV are shown in orange, and contacts unique to SFV are shown in beige. **(E)** Staining of HEK293T cells transfected to express wild-type or mutant SFV E3–E2–(6K/TF)–E1 proteins by the indicated Fc fusion proteins. See Methods and Figure S9B for additional information. **(F)** Ribbon and surface diagram showing the contacts between SFV E2–E1 glycoproteins and VLDLR (PDB: 8UA8) (*5*) **(G–I)** K562 cells ectopically overexpressing *Hs*MXRA8, *Ae. albopictus* Lachesin, or *Ae. aegypti* LpR1 were infected with GFP-expressing wild-type SFV (strain SFV4) RVPs (**G**), E2 157A–160A mutant SFV RVPs (**H**), or E1 K345A and K347A mutant SFV RVPs (**I**) at an MOI of 2 (measured on K562 cells expressing Ae. albopictus Lachesin) (**G, H**) or at an MOI of 0.5 (measured on K562 cells expressing *Ae. aegypti* LpR1) (**I**). Infectivity was measured using flow cytometry. Data are mean ± s.d. from three experiments performed in duplicate (*n* = 3) (**G**–**I**). One-way ANOVA with Dunnett’s multiple comparisons test, \*\*\*\**P* < 0.0001 (**G**–**I**).

We next transfected HEK293T cells to express WT or mutant SFV E3–E2–(6K/TF)–E1 proteins containing alanine substitutions of E2 residues whose side chains contact Lachesin and assessed cell staining by Lachesin–Fc (Fig. S9B). The E2 Q157A–T160A quadruple alanine mutant, which we hypothesized would disrupt SFV E2 β-ribbon connector central arch contacts with Lachesin (Fig. S11C), abolished Lachesin–Fc staining (Fig. 2E), indicating that this E2 segment is critical for Lachesin engagement by SFV and CHIKV. However, among these four residues, the most important E2 β-ribbon contributor to Lachesin binding differs between SFV and CHIKV. Alanine substitution of CHIKV E2 Q158 alone almost completely abolished Lachesin–Fc staining (Fig. 1G), but the equivalent SFV E2 Q158A substitution only had a minor effect on staining (Fig. 2E). In contrast, while the CHIKV E2 S159A substitution had no effect on Lachesin–Fc staining (Fig. 1G), the SFV E2 T159A substitution markedly diminished Lachesin–Fc staining (Fig. 2E).

Alanine substitutions of SFV E2 residues R144, T264, and R266, which make polar contacts with the N-terminal region of Lachesin D1 and loop C–D (Fig. S11D), decreased Lachesin–Fc immunostaining (Fig. 2E). Other alanine substitutions had minimal to no effect, suggesting that the large SFV-Lachesin complex interface can tolerate loss of most individual contact residues.

SFV can bind low-density lipoprotein receptor-related proteins including VLDLR and ApoER2 (also known as LRP8) to infect vertebrate cells and can additionally recognize Lipophorin receptor 1 (LpR1), which is a mosquito ortholog of VLDLR (*26-28*). The ectodomain of VLDLR contains a cell membrane-distal ligand-binding domain (LBD), which comprises eight LDLR class A (LA) repeats. LA repeats found in VLDLR and ApoER2 bind domain III (DIII) in the SFV E1 glycoprotein near the twofold and fivefold symmetry axes (Fig. 2F) (*5, 34-37*).

Given that VLDLR and Lachesin bind completely distinct surfaces on the SFV E2–E1, we next sought to confirm that interactions with LA repeats and Lachesin could be individually disrupted. VLDLR LA repeats use conserved aspartic acid and tryptophan residues to contact K345 and K347 in SFV E1 DIII (Fig. 2F) (*5, 34*). In HEK293T cells transfected to express WT and mutant SFV E2–E1, alanine substitutions of SFV E1 residues K345 and K347 diminished staining by an Fc fusion protein containing the human VLDLR LBD (VLDLR_LBD_–Fc) but had no effect on Lachesin–Fc staining (Fig. 2E). Additionally, all tested alanine substitutions of Lachesin-contacting residues had little to no effect on human VLDLR_LBD_–Fc staining (Fig. 2E).

SFV RVPs bearing WT E2–E1, as expected based on prior studies, infected K562 cells expressing *Ae. albopictus* Lachesin or *Ae. aegypti* LpR1 (chosen because of better expression as compared to *Ae. albopictus* LpR1 when expressed in K562 cells) (*1, 27*) (Figs. 2G and S7D). SFV RVPs containing the K345A and K347A E1 substitutions, however, did not infect cells expressing LpR1 but retained the ability to infect cells expressing Lachesin (Fig. 2H). Conversely, SFV RVPs containing alanine substitutions of E2 β-ribbon connector residues 157–160, which impair Lachesin–Fc recognition, could not infect cells expressing Lachesin, but could still infect cells expressing LpR1 (Fig. 2I). These data confirm that SFV E2–E1 LA repeat and Lachesin binding sites are structurally and functionally separate.

### MIDV uses an altered Lachesin binding mode

MIDV is an alphavirus that is genetically related to the SF complex but is grouped in a different antigenic complex, the MID complex (Fig. 1A). CHIKV and MIDV E2 only share 52% amino acid sequence identity (Fig. 2A). While CHIKV and SFV E2 β-ribbon residues 157– 159 contain determinants of Lachesin binding, these residues are divergent in the MIDV E2 sequence: residues “VQS” and “QQT” found in CHIKV and SFV, respectively, are replaced with “AMR” (Fig. S12). To understand how MIDV binds Lachesin despite sequence divergence in E2 β-ribbon central arch residues, we obtained a cryo-EM structure of MIDV VLPs in complex with Lachesin–Fc, with focused refinement of the asymmetric unit yielding a 2.9 Å reconstruction (Figs. 3A, 3B, S13, and Table S1).

**Figure 3.**
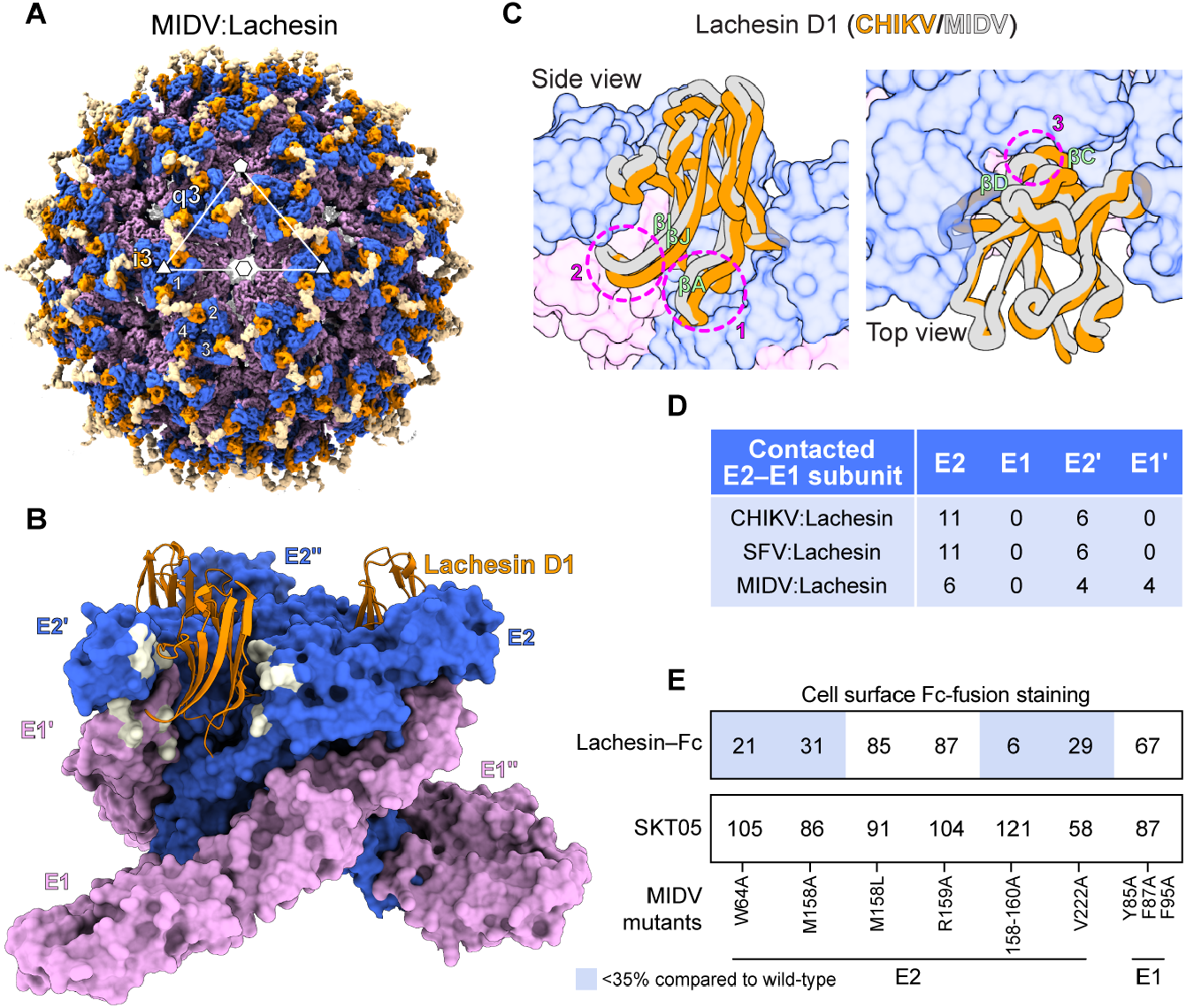
MIDV recognition of *Aedes albopictus* Lachesin. **(A)** Overall cryo-EM map of Lachesin–Fc bound to MIDV VLP. Density for the MIDV E2 and E1 glycoproteins are shown in blue and pink, respectively. Density for bound Lachesin is colored orange for D1 and beige for additional density. Icosahedral symmetry axes (i5, i3, i2) are indicated with a pentagon, triangle, and hexagon, respectively. The icosahedral i3 and q3 E2–E1 trimers are indicated. Four E2–E1 heterodimers that constitute an asymmetric unit are numbered. **(B)** MIDV E2–E1 trimer bound to Lachesin D1. The MIDV E2–E1 trimer is surface rendered, and Lachesin D1 molecules are shown as ribbon diagrams. MIDV E2–E1 residues that interact with D1 are shown in yellow. See Figure S14 for detailed interactions. **(C)** Superposition of CHIKV E2–E1 and MIDV E2–E1 bound to Lachesin D1, aligned based on E2–E1 residues. A surface representation of E2 and E1 is shown in blue and pink, respectively. Circled regions with associated numbers indicate regions with conformational changes in Lachesin D1 between CHIKV and MIDV structures. 1: rearrangement of the N-terminal A β-strand, 2: repositioning of the I–J loop, 3: shifting of the C–D loop. **(D)** Summary number of Lachesin contact residues on the CHIKV, SFV, and MIDV E2–E1 glycoproteins and the adjacent protomer (E2′–E1′). **(E)** Staining by the indicated Fc fusion proteins of cells transfected to express wild-type or mutant E3–E2– (6K/TF)–E1 proteins of MIDV. Staining with a humanized anti-E1 cross-reactive antibody SKT05 (hSKT05) (*4*) was used to normalize Fc-fusion protein staining. Data are mean from three experiments (*n* = 3). See Methods and Figure S9C for additional information.

Structural comparison of the MIDV- and CHIKV-Lachesin complexes revealed remodeled receptor-viral glycoprotein contacts caused by conformational rearrangements in three Lachesin D1 regions (Figs. 3C and S14A–E). These differences in binding mode cause the majority of Lachesin contacts to be redistributed from the E2–E1 protomer to the neighboring E2’–E1’ protomer (Fig. 3D). Lachesin maintains generally similar interactions with E2 domain A (Fig. S14B) and E2’ domain B (Fig. S14E) as observed in CHIKV and SFV, and new MIDV-specific contacts with E1’ are established that include interactions with conserved fusion loop residues F87 and F95, which are also contacted by MXRA8 (Figs. S14D and S15). While there are similarities in how Lachesin interacts with the central arch of the E2 β-ribbon connector in the CHIKV and SFV receptor-bound structures, there are major differences in how Lachesin interacts with the MIDV E2 β-ribbon connector (Figs. S3C, S11C, and S14C). Rather than the receptor-viral glycoprotein interface at this site being dominated by side chain contacts, most interactions with the β-ribbon connector involve backbone atoms.

We ectopically expressed WT or mutant MIDV glycoproteins containing alanine substitutions of Lachesin-contacting residues in HEK293T cells and stained cells with Lachesin–Fc (Figs. 3E and S9C). Simultaneous alanine substitution of MIDV E2 β-ribbon connector residues M158– T160 abolished Lachesin–Fc staining (Fig. 3E). Even though the M158 side chain points away from Lachesin (Fig. S14C), the E2 M158A mutation markedly impaired Lachesin–Fc staining (Fig. 3E). The side chain of M158 is in a cluster of hydrophobic residues in E2 domain A and the E2 β-ribbon connector (Fig. S14F). We thus hypothesized that M158 anchors the conformation of the polypeptide backbone of the central arch of the E2 β-ribbon connector to allow for productive Lachesin engagement. Consistent with this hypothesis, MIDV E2–E1 containing the M158L substitution, which would preserve contacts with the hydrophobic cluster of residues, bound Lachesin–Fc (Fig. 3E).

Multiple other alanine substitutions of Lachesin-contacting residues in MIDV differ in their effects on Lachesin–Fc staining from their CHIKV and SFV counterparts; alanine substitutions at E2 residues W/Y64 or E2’ I/V222 had little to no effect on CHIKV and SFV Lachesin binding but profound effects on MIDV Lachesin binding (Figs. 1G, 2E, and 3E). Although MIDV makes unique contacts with the E1’ fusion loop (including residues F87 and F95), simultaneous alanine substitution of multiple fusion loop aromatic residues had little effect on Lachesin– Fc staining (Figs. 3E and S14D). Thus, despite some differences in specific residue contacts and binding modes among CHIKV, SFV, and MIDV, the central arch of the E2 β-ribbon connector plays a critical role in Lachesin recognition by all tested arthritogenic alphaviruses.

### Lachesin determinants of alphavirus engagement

The E2–E1 glycoproteins of the arthritogenic alphaviruses MAYV, RRV, and ONNV can bind *Ae. albopictus* Lachesin (*1*). We next performed RVP infectivity assays on K562 cells ectopically expressing unmutated or mutant *Ae. albopictus* Lachesin containing an internal Flag tag (Figs. S7E and S7F). Lachesin residues F49 and S70 are on opposite faces of the cleft in structures of Lachesin-bound CHIKV E2–E1 glycoproteins (Fig. 4A). If the general Lachesin binding mode is shared among arthritogenic alphaviruses, arginine substitution of Lachesin F49 or S70 would introduce steric clashes with alphavirus E2 glycoproteins and disrupt binding by MAYV, RRV, and ONNV, alphaviruses for which we have not determined Lachesin-bound structures. The S70R mutation completely abolished CHIKV, SFV, MIDV, MAYV, RRV, and ONNV RVP infection of K562 cells expressing the mutant receptor (Fig. 4B). The F49R mutation completely abrogated entry for most tested alphavirus RVPs except for MAYV and RRV, for which the phenotype was less severe (Fig. 4B). These findings are consistent with a generally similar binding mode of these alphaviruses with Lachesin.

**Figure 4.**
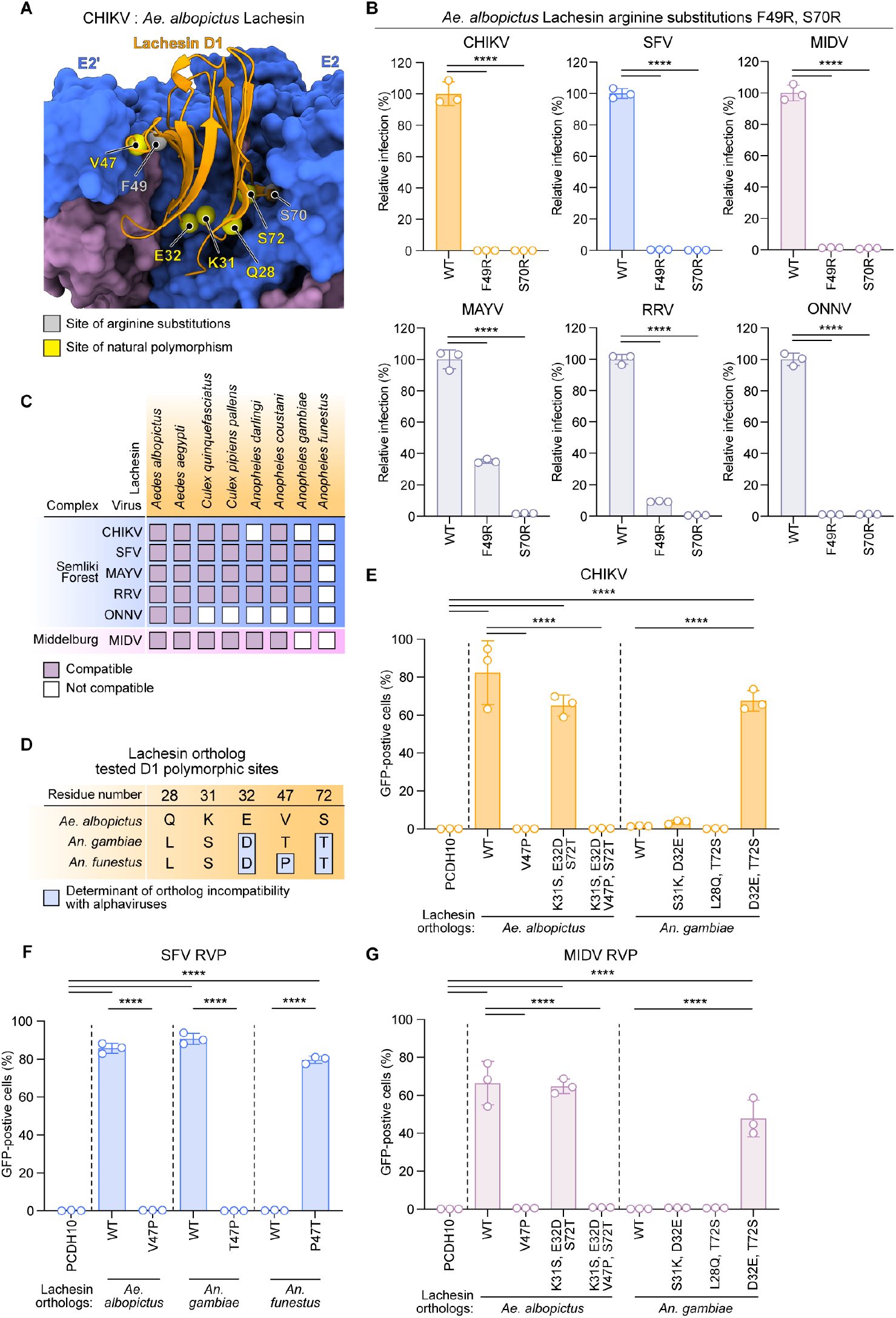
Lachesin determinants of alphavirus binding. **(A)** Surface representation of CHIKV E2–E1 and ribbon diagram of *Ae. albopictus* Lachesin D1. Spheres indicate the positions of Lachesin residues subjected to mutational analysis. Gray spheres are residues mutated to arginine to introduce steric clashes. Yellow spheres represent select polymorphic residues between the *Ae. albopictus, An. gambiae*, and *An. funestus* orthologs. **(B)** Infection of K562 cells expressing wild-type (Lachesin–Flag) or mutant Lachesin constructs containing the indicated arginine substitutions by GFP-expressing alphavirus RVPs for CHIKV 37997, SFV SFV4, MIDV MIDV857, MAYV LET-1430, RRV T48, or ONNV SG650 RVPs at MOI of 2 (measured on K562 cells expressing *Ae. albopictus* Lachesin). Infection was quantified using flow cytometry. **(C)** Summary of mosquito Lachesin ortholog usage by CHIKV, SFV, MAYV, RRV, ONNV, and MIDV (*1*). **(D)** Summary of polymorphic Lachesin D1 residues that determine the inability of *An. gambiae* or *An. funestus* Lachesin orthologs to support CHIKV or SFV E2–E1-dependent entry. **(E–G)**, Infection of K562 cells expressing wild–type or mutant *Ae. albopictus, An. gambiae* Lachesin orthologs by GFP-expressing CHIKV 37997 (**E**), SFV SFV4 (**F**), MIDV MIDV857 (**G**), at MOI of 1 (measured on K562 cells expressing *Ae. albopictus* Lachesin). Infectivity was quantified using flow cytometry. Data are mean ± s.d. from three experiments performed in duplicates (*n* = 3) (**B, E, F, G**). One-way ANOVA with Dunnett’s multiple comparisons test, \*\*\*\**P* < 0.0001 (**B**). One-way ANOVA with Šídák’s multiple comparisons test, \*\*\*\**P* < 0.0001 (**E, F, G**).

Arthritogenic alphaviruses broadly recognize multiple, but not all mosquito orthologs of Lachesin (*1*) (Fig. 4C). For example, CHIKV cannot infect cells that express the Lachesin ortholog of *An. gambiae*, which is a vector for ONNV and malaria. Additionally, the Lachesin ortholog of *An. funestus*, another ONNV and malaria vector, is not recognized by any of the tested alphaviruses. Most of the *Ae. albopictus* Lachesin residues that CHIKV, SFV, and MIDV contact are conserved across orthologs of *Aedes, Anopheles*, and *Culex* mosquito species (Fig. S16A), suggesting that small numbers of natural polymorphisms in Lachesin ortholog amino acid sequences likely underlie the incompatibility of certain *Anopheles* Lachesin orthologs with alphaviruses.

Examination of Lachesin D1 sequence alignments and the cryo-EM structures revealed that, with respect to *Ae. albopictus* Lachesin, *An. funestus* Lachesin contains a V47P polymorphism that may alter interactions a Lachesin loop makes with E2’ domain B in the receptor-bound CHIKV, SFV, and MIDV structures (Figs. 4D, S3E, S11E, and S14E). Indeed, a single V47P substitution in *Ae. albopictus* Lachesin abolished CHIKV, SFV, and MIDV RVP entry into K562 cells (Figs. 4E–G and S7G). Three other substitutions in *Ae. albopictus* Lachesin with *An. funestus* polymorphic residues that either remove contacts with CHIKV E2 or introduce steric clashes (K31S, E32D, S72T) had little effect on CHIKV RVP entry even when combined (Fig. 4E, S3C, and S3D).

SFV can bind *An. gambiae* Lachesin, which, with respect to *Ae. albopictus* Lachesin, contains the V47T substitution (Fig. 4D). *An. gambiae* Lachesin with the single T47P substitution did not support SFV RVP entry; conversely, introducing the P47T substitution in *An. funestus* Lachesin allowed it to mediate SFV RVP entry (Fig. 4F). When tested for its impact on the entry mediated by the E2–E1 proteins of other arthritogenic alphaviruses for which we did not obtain cryo-EM structures, we found that the presence of a proline at Lachesin residue 47 critically modulated the entry of RRV, MAYV, and ONNV RVPs (Fig. S16B–D).

*An. gambiae* Lachesin, which contains T47, can serve as a receptor for SFV but not CHIKV or MIDV (Fig. S16D). We next tested whether substituting residues in *An. gambiae* Lachesin with residues that are present in *Ae. albopictus* Lachesin would allow the modified *An. gambiae* Lachesin to serve as a CHIKV receptor. The D32E and T72S substitutions when jointly introduced, but not the other double mutants, converted *An. gambiae* Lachesin into a compatible receptor for CHIKV and MIDV (Figs. 4E and 4G). Therefore, the inability of *An. gambiae* Lachesin to support CHIKV E2–E1-dependent entry is likely determined by the combined effects of multiple unfavorable polymorphic residues in the contact interface.

Together, these results suggest that *An. funestus* Lachesin residue P47 is a critical determinant of the inability of that ortholog to serve as a receptor for CHIKV and almost all other tested arthritogenic alphaviruses.

### Lachesin–Fc protects mice against alphavirus pathogenesis in vivo

Soluble receptor–Fc decoy proteins can inhibit alphavirus infection in cell culture and confer protection in animal models of alphavirus challenge (*2, 38, 39*). We tested Lachesin–Fc and *Hs*MXRA8–Fc decoy proteins for their ability to neutralize CHIKV, SFV, RRV, GETV, MAYV, and MIDV RVP entry into Vero E6 cells at a single concentration of 100 µg ml^-1^ (Fig. 5A). Lachesin–Fc robustly neutralized the entry of all tested RVPs, indicating broad activity (Fig. 5B). An Fc fusion protein containing the EC1 repeat of human PCDH10 (*Hs*PCDH10_EC1_–Fc, Fig. S2E), used as a control, had no activity. In contrast, *Hs*MXRA8–Fc strongly neutralized CHIKV RVP entry, had partial neutralizing activity against MAYV and RRV RVPs, but did not neutralize SFV or GETV RVP entry.

**Figure 5.**
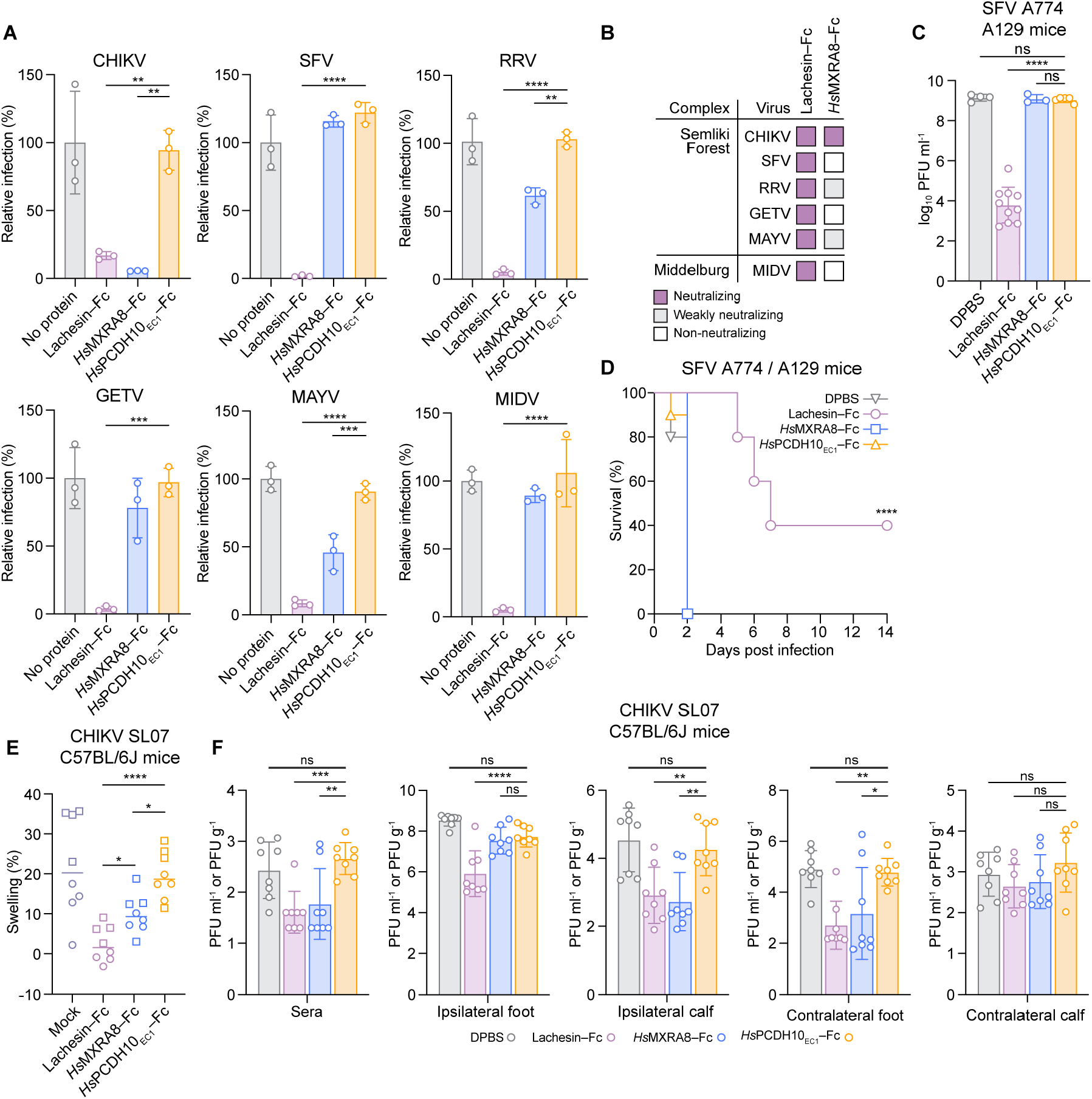
Lachesin–Fc receptor decoy broadly neutralizes arthritogenic alphaviruses and is protective in vivo. **(A)** Neutralization of GFP-expressing RVPs for the indicated alphaviruses by Lachesin–Fc, *Hs*MXRA8–Fc, and *Hs*PCDH10_EC1_–Fc in Vero cells at an MOI of 1 (measured on K562 cells expressing Lachesin) and Fc fusion protein concentration of 100 µg ml^-1^. Infection was monitored using flow cytometry. RVP E2–E1 sequences are from strains CHIKV 37997, SFV SFV4, RRV T48, GETV M1, MAYV LET-1430, and MIDV MIDV857. **(B)** Summary of neutralization assays performed in (**A**). “Neutralizing” signifies greater than 90% neutralization at the tested concentration. “Weakly neutralizing” signifies less than 75% neutralization at the tested concentration. **(C**,**D)** Six-to eight-week-old A129 mice were administered *Ae. albopictus* Lachesin–Fc, *Hs*MXRA8– Fc fusion protein, *Hs*PCDH10_EC1_–Fc (isotype control), or a DPBS buffer (mock) intraperitoneally 6 h before subcutaneous inoculation with 10^4^ PFU of SFV A774 rescued from a molecular clone. At two days post-infection, SFV was measured in sera (**C**). Survival of the mice was monitored against SFV challenge (**D**). Group sizes were: DPBS only, *n* = 10; Lachesin–Fc, *n* = 10; *Hs*MXRA8–Fc, *n* = 10; *Hs*PCDH10_EC1_–Fc, *n* = 10 mice. **(E**,**F)** Four-week-old mixed-sex C57BL/6J mice (*n* = 8 per group; 4 males and 4 females) were treated intraperitoneally with Lachesin–Fc, *Hs*MXRA8–Fc, *Hs*PCDH10_EC1_–Fc, or an equivalent volume of DPBS 24 h before subcutaneous footpad inoculation with CHIKV strain SL07. Footpad thickness was measured immediately before CHIKV strain SL07 infection and again at 3 days post infection (dpi) to assess swelling (**E**). At 3 dpi, CHIKV titers were quantified in sera, ipsilateral foot and calf, and contralateral foot and calf (**F**). Data are mean ± s.d. from three experiments performed in duplicate (*n* = 3) (**A**). One-way ANOVA with Dunnett’s multiple comparisons test (**A, C, F**). \*\**P =* 0.0048 (CHIKV Lachesin–Fc) \*\**P =* 0.0021 (CHIKV *Hs*MXRA8– Fc) **P = 0.0019 (RRV HsMXRA8–Fc) \*\*\**P =* 0.004 (GETV Lachesin–Fc) \*\*\**P* = 0.0007 (MAYV *Hs*MXRA8–Fc) \*\*\*\**P* < 0.0001 (**A**). \*\*\*\**P <* 0.000001 (**C**). Log-rank (Mantel-Cox) test \*\*\*\**P* = 0.000021 (**D**). Brown–Forsythe and one-way Welch ANOVA with Dunnett T3 multiple comparisons test \**P =* 0.017631 (*Hs*MXRA8–Fc and Lachesin–Fc) \**P =* 0.017631 (*Hs*PCDH10_EC1_–Fc and *Hs*MXRA8–Fc) \*\*\*\**P* < 0.0001 (**E**) \**P* = 0.018809 (Contralateral foot *Hs*MXRA8–Fc) \*\**P =* 0.004750 (Sera *Hs*MXRA8–Fc) \*\**P =* 0.009292 (Ipsilateral calf Lachesin–Fc) \*\**P* = 0.003102 (Ipsilateral calf *Hs*MXRA8– Fc) \*\**P* = 0.002379 (Contralateral foot Lachesin–Fc) \*\*\**P* = 0.000922 (Sera Lachesin–Fc) \*\*\*\**P* < 0.0001 (**F**)

A129 mice are deficient in type I interferon signaling and experience lethal outcomes to alphavirus infection, offering a stringent model to evaluate protective activity of anti-alphavirus countermeasures in vivo (*40*). While all A129 mice that received Dulbecco’s phosphate-buffered saline (DPBS, vehicle), *Hs*PCDH10_EC1_–Fc (control protein), or *Hs*MXRA8–Fc succumbed to infection within 48 h of challenge with a lethal dose (10^4^ PFU) of SFV, Lachesin–Fc-treated animals experienced a five-log decrease in viremia and 40% of mice survived the challenge (Figs. 5C and 5D).

C57BL/6J mice are an established model of arthritogenic alphavirus challenge as they experience nonlethal infection but develop arthritic signs (*41*). Mice were administered Lachesin–Fc, *Hs*MXRA8–Fc, *Hs*PCDH10_EC1_–Fc as a control, or DPBS and were challenged 24 h later with CHIKV through footpad inoculation. Both Lachesin–Fc and *Hs*MXRA8–Fc reduced foot swelling, with Lachesin–Fc conferring more pronounced reduction than *Hs*MXRA8–Fc (Fig. 5E). Additionally, three days postinfection, mice that received Lachesin–Fc or *Hs*MXRA8–Fc showed reduced viral RNA levels in serum, ipsilateral calf and foot, and contralateral foot (Fig. 5F). Together, these data establish Lachesin–Fc as an efficacious decoy receptor against multiple alphaviruses in the SF and MID complexes with demonstrated in vivo efficacy and broader activity than MXRA8–Fc.

## Discussion

Gene drive is a naturally occurring form of super-Mendelian genetic element that can be harnessed to modify alleles of wild insect populations (*42*). We identified a single polymorphism (V47P) in the Lachesin ortholog of *An. funestus*, also a malaria and ONNV vector, that is sufficient to prevent CHIKV, SFV, and MIDV E2–E1 engagement when introduced into *Ae. albopictus* Lachesin. The V47P mutation is thus an attractive candidate for engineering resistant mosquito vectors. Because this mutation is inspired by a natural polymorphism found in *An. funestus* mosquitoes, we expect it to preserve the native, cell-adhesion function of Lachesin and mosquito fitness, selectively ablating virus entry with minimal fitness cost to the mosquito vector, which may be an important component of certain ecosystems.

MAYV was the only tested alphavirus for which E2–E1 could still recognize *Ae. albopictus* Lachesin containing the V47P substitution (Fig. S16B). However, P47 nonetheless likely impairs MAYV E2–E1 binding, given that MAYV could efficiently recognize *An. funestus* Lachesin containing the P47T substitution, despite not recognizing unmutated *An. funestus* Lachesin. These findings suggest that *An. funestus* Lachesin contains determinants other than P47 that account for its inability to be recognized by MAYV.

Interestingly, MAYV RVP entry was the least impacted by the F49R substitution, which, like the V47P substitution, would disrupt contacts Lachesin makes with E2′ domain B (Figs. 4A and 4B). These findings suggest the contacts Lachesin makes with MAYV E2′ domain B may be altered as compared to CHIKV, SFV, and MIDV.

CHIKV binds both a mammalian receptor, MXRA8, and a mosquito receptor, Lachesin, in clefts formed by adjacent E2–E1 protomers, which is the most common receptor-binding surface for alphaviruses (*5, 25, 31, 43-46*). The cleft provides a large interaction surface to accommodate receptor-specific contacts. Our analyses suggest that contact residues that contribute most prominently to CHIKV’s affinities for a human receptor, MXRA8, and a mosquito receptor, Lachesin, are functionally separate, a conclusion that is also supported by DMS mapping of human versus mosquito determinants of CHIKV cellular entry (*33*).

The ability of the CHIKV E2–E1 cleft to bind multiple receptors is reminiscent of that of the encephalitic alphavirus WEEV, which uses the same cleft to accommodate two avian receptors, MXRA8 and PCDH10, in addition to VLDLR, through distinct binding modes (*31, 45-49*). In WEEV, the E2–E1 glycoprotein residues that most strongly contribute to binding to each of these multiple receptors are also largely separate (*31, 46*). Use of different residues within a shared receptor binding surface as the principal determinants for engagement of distinct receptors may represent a common structural arrangement among alphaviruses to maintain fitness in diverse hosts.

SFV uses VLDLR, not MXRA8, as a receptor on vertebrate cells (*27*). While the cleft appears to be a canonical receptor-binding surface for alphaviruses, SFV binds LA repeats at a site on E1 DIII that does not overlap with the Lachesin interface (*5, 34, 37*), and seems to use this same site to bind LpR1, a mosquito ortholog of VLDLR (Figs. 2F and 2H). SFV can thus use distinct surfaces to engage Lachesin and LpR1 as alternative receptors on mosquito cells.

The CHIKV E1 A226/V226 polymorphism defines some strains of the Indian Ocean sublineage in the East/Central/South Africa lineage of CHIKV and has been shown to enhance CHIKV fitness in *Ae. albopictus* mosquitoes (*50*). As *Ae. albopictus* Lachesin contacts CHIKV E2 and makes no E1 contacts, the structural studies presented here corroborate previous work suggesting that the E1 A226V polymorphism acts downstream of cellular receptor binding (*50, 51*).

While receptor decoy proteins have shown promise as antivirals for alphaviruses, MXRA8–Fc, a previously studied decoy receptor against arthritogenic alphaviruses (*2*), has limited breadth. We show that Lachesin–Fc has robust neutralizing activity against MAYV, RRV, and GETV, in addition to neutralizing SFV and MIDV, which are two viruses that do not bind MXRA8. Lachesin receptor decoys are thus promising antiviral agents with enhanced breadth for epidemic preparedness.

## Acknowledgements

This work was supported by a Burroughs Wellcome PATH award to J.A., a Vallee Scholar award to J.A., a Smith Family Foundation Odyssey award to J.A., a Charles E.W. Grinnell Trust award to J.A., NIH award R01 AI182377 to J.A., and a G. Harold and Leila Y. Mathers Foundation award to J.A. This work was also supported by the UTMB Institute for Human Infections and Immunity award to K.S.P. and by NIH awards T32AI700245 to J.S.P., and R24AI120942 to S.C.W. J.A. is an investigator of the Howard Hughes Medical Institute. We thank Bridget Golan for assistance with illustration editing. This article is subject to the HHMI Open Access to Publications policy. HHMI lab heads have previously granted a nonexclusive CC BY 4.0 license to the public and a sublicensable license to HHMI in their research articles. Pursuant to those licenses, the author-accepted manuscript of this article can be made freely available under a CC BY 4.0 license immediately upon publication.

## Author contributions

X.F. performed cryo-electron microscopy and built the models, W.L. performed glycoprotein cell surface staining experiments with assistance from J.S.P. C.E.H. and C.J. helped with construct generation and cell-surface immunostaining experiments. W.L. and J.S.P. purified Fc fusion proteins. J.S.P. performed infectivity studies with assistance from W.L. J.A.P. designed and conducted in vivo studies using replication-competent SFV and CHIKV with the help of E.M.H. and N.C.H.; S.C.W. and K.S.P. supervised those experiments. S.C.W., K.S.P., and J.A. acquired funding. X.F., W.L., and J.S.P. wrote the original draft of the manuscript, and all authors participated in reviewing and editing.

## Materials and Methods

### Cell lines and viruses

HEK293T (human kidney epithelial, ATCC CRL-11268) and Vero E6 (*Cercopithecus aethiops* kidney epithelial, ATCC CRL-1586) were maintained in Dulbecco’s modified Eagle’s medium (DMEM, Gibco) supplemented with 10% (v/v) fetal bovine serum (FBS) and 25 mM HEPES (Thermo Fisher Scientific). K562 (human chronic myelogenous leukemia, ATCC CCL-243) cells were maintained in RPMI-1640 (Thermo Fisher Scientific) supplemented with 10% (v/v) FBS, 25 mM HEPES, and 1% (v/v) penicillin-streptomycin. BHK cells were grown in similar media with the addition of 1× GlutaMAX (Gibco), 1× MEM non-essential amino acids (Gibco), and 1 mM sodium pyruvate (Gibco). Both cell types were maintained at 37°C with 5% CO2. Expi293F cells (Thermo Fisher Scientific A14527) were maintained in Expi293 Expression Medium (Thermo Fisher Scientific). Cell lines were not authenticated. Absence of mycoplasma was confirmed through monthly mycoplasma tests using the e-Myco PCR detection kit (Bulldog Bio 25234).

Chikungunya virus (strain SL07) and Semliki Forest virus (strain A774) were rescued from infectious clones. To recover chikungunya (CHIKV) SL07 (*52*), RNA from 1 µg linearized plasmid was in vitro transcribed with the SP6 mMessage mMachine kit (Invitrogen, Carlsbad, CA). The RNA was electroporated into 10^7^ BHK cells in a 2 mm gap cuvette using an ECM 830 (Harvard Apparatus, Holliston, MA) delivering eight 100 µs pulses of 650 volts with intervals of 300 µs. A plasmid encoding infectious cDNA of SFV A774wt (a gift from A. Merits) (*53*) was electroporated into BHK-21 cells. Cells were then plated in a 100-mm dish and incubated at 37 °C for 24 h to obtain a P0 stock. To obtain a P1 stock of the virus, BHK-21 cells were infected with P0 virus at a multiplicity of infection (MOI) of 5. Resulting P1 virus was harvested 24 h post infection. Titers for SFV strain A774 and CHIKV strain SL07 were determined by plaque assay on Vero E6 cells.

### Production of virus-like particles

Plasmids encoding the structural polyprotein (capsid–E3–E2–[6K/TF]–E1) of CHIKV strain 37997 (*54*) (GenBank: AAU43881) and MIDV strain MIDV857 (GenBank: EF536323), were transfected into Expi293F™ cells using the ExpiFectamine^TM^ 293 Transfection Kit (Thermo Fisher Scientific A14525) according to the manufacturer’s protocol. Culture supernatants were collected 5 d post-transfection and clarified by centrifugation at 3,000 × g for 20 min. Clarified supernatants were layered onto 35% (w/v) and 70% (w/v) sucrose cushions and ultracentrifuged at 150,000 × g for 3 h in a Beckman SW32Ti rotor at 4 °C. VLPs were harvested from the 35%/70% interface, loaded onto a 20–70% (w/v) continuous sucrose gradient, and further purified by ultracentrifugation at 210,000 × g for 1.5 h in a Beckman SW41 rotor at 4 °C. EEEV VLPs were generated in a similar manner using a vector encoding the structural polyprotein of EEEV strain PE6 (GenBank: AAU95735.1) containing the K67N capsid mutation used to improve the yield of VLP production. (*55*)

SFV VLPs were produced by transiently transfecting adherent HEK293T cells with a plasmid encoding the SFV4 structural polyprotein (*5*) (GenBank: AKC01668.1) using Lipofectamine 3000 (Invitrogen), following the manufacturer’s instructions. Culture supernatants were collected 48 h post-transfection and clarified by centrifugation at 3,000 × g for 20 min. VLPs were pelleted through a 30% (w/v) sucrose cushion at 110,000 × g for 2.5 h in a Beckman SW32Ti rotor at 4 °C, resuspended in DPBS (Thermo Fisher Scientific 14190-144), and further purified on a 20–70% (w/v) sucrose gradient at 210,000 × g for 1.5 h in a Beckman SW41 rotor at 4 °C. The VLP band was collected and stored at 4 °C without freezing. Particle integrity and the absence of degradation products were confirmed by SDS-PAGE. VLP preparations were used within 7 days of purification, with buffer exchange performed immediately before use as required by the downstream assay.

### Expression and purification of recombinant proteins

The genes encoding *Aedes albopictus* Lachesin (residues 1–345; GenBank: XP_019932516.3), the human MXRA8 ectodomain (residues 20–337; GenBank: NP_001269511.1) (*27*), the mouse MXRA8 ectodomain (residues 21-340; GenBank: NP_077225.4), and the EC1 domain of human PCDH10 (residues 19–122; GenBank: NP_116586.1) (*31*) were each cloned into the pVRC expression vector with a human IgG1 Fc tag fused at the C terminus.

For production of Lachesin–Fc, *Hs*PCDH10EC1–Fc, and *Hs* and *Mm*MXRA8–Fc, Expi293F™ cells were transiently transfected with the corresponding expression plasmids using the ExpiFectamine™ 293 Transfection Kit according to the manufacturer’s instructions. Supernatants were harvested 5 d post-transfection, clarified by centrifugation at 4,000 × g for 30 min, and applied to MabSelect™ PrismA protein A resin (Cytiva 17549801) for affinity purification following the manufacturer’s protocol. The proteins were further purified by size-exclusion chromatography on a Superdex 200 Increase 10/300 column (Cytiva) and stored in Tris-buffered saline (20 mM Tris, 150 mM NaCl, pH 7.5). For in vivo studies, proteins were dialyzed immediately into DPBS buffer after elution and not subjected to size-exclusion chromatography, except for a small aliquot analyzed for quality control purposes, which confirmed that the material eluted as a single peak. Endotoxin levels were quantified using a Pierce Chromogenic Endotoxin Quantification Kit (Thermo Fisher Scientific A39553) and were <0.5 endotoxin units mg^−1^.

### Biolayer interferometry binding assays

Biolayer interferometry was performed on an Octet RED96e instrument (Sartorius), and data were analyzed using ForteBio Data Analysis HT software (version 12.0.1.55). The anti-CHIKV E2–E1 monoclonal antibody Chk265 (absolute antibody Ab00687-1.1) was immobilized onto Anti-Mouse IgG Fc Capture (AMC) biosensors (Sartorius 18-5088) at 250 nM in kinetic buffer (TBS supplemented with 2 mM CaCl_2_, 0.1% (w/v) bovine serum albumin, and 0.01% Tween) for 600 s. After a 60 s baseline in kinetic buffer, sensors were transferred to wells containing 1 µM CHIKV VLPs for 1 h, followed by an additional 1 h equilibration in kinetic buffer. Sensors were then washed in kinetic buffer for 60 s to re-establish a baseline and moved into wells containing serial dilutions of Lachesin–Fc (100–6.25 nM in kinetic buffer) for a 300 s association phase, followed by a 300 s dissociation phase in kinetic buffer. Binding kinetics were determined using a standard 1:1 interaction model.

### ELISA

Apparent affinities of Lachesin–Fc, *Hs*MXRA8–Fc, and *Hs*PCDH10_EC1_–Fc for CHIKV, SFV, and MIDV VLPs were determined by ELISA. Briefly, 200 ng of VLPs were immobilized on ELISA MaxiSorp plates (Thermo Scientific 439454) overnight at 4 °C. Plates were blocked with PBS containing 3% (w/v) bovine serum albumin (BSA) for 1 h at room temperature and washed three times with PBS. Serial dilutions of Lachesin–Fc, *Hs*MXRA8–Fc, or *Hs*PCDH10_EC1_–Fc were added and incubated for 1 h at room temperature. After three washes with PBS, 100 µl per well of horseradish peroxidase– conjugated anti-human IgG (Sigma A0170), diluted 1:20,000 in PBS with 3% BSA, were added and incubated for 1 h at room temperature. Plates were washed five times with PBS, developed with 100 µl per well of 1-step TMB ELISA substrate for 3 min at room temperature in the dark, and reactions were stopped with 100 µl per well of 2 N sulfuric acid. Absorbance was measured at 450 nm using a BioTek multimode reader. EEEV PE6 VLPs were included as negative controls.

### Truncation and mutagenesis, and stable cell line generation

K562 cells stably transduced to express *Aedes albopictus* Lachesin (GenBank XP_019932516.3), human MXRA8 (GenBank NM_032348.3), human VLDLR (GenBank NP_003374), N-term Flag murine MXRA8 (GenBank XP_006539298), and *Ae. aegypti* LpR1 (GenBank AEY84776) were generated in prior studies and cell surface staining was again confirmed here.

For the K562 cells ectopically expressing Flag-tagged wild-type or mutant *Ae. albopictus* Lachesin, we ordered cDNA encoding human codon-optimized *Ae. albopictus* Lachesin (GenBank XP_019932516) from Integrated DNA Technologies (IDT) containing an internal FLAG in the stalk region after D3 (between V308 and I309). Single and double amino acid mutations were introduced using primers encoding mutations.

The above constructs were cloned into lentiGuide-Puro (a gift from F. Zhang; Addgene #52963) (*56*) and transfected with psPAX2 (a gift from D. Trono; Addgene # 12260) and pMD2.G (a gift from D. Trono; Addgene # 12259) at a ratio of 3:2:1 into HEK293T cells using Lipofectamine^TM^ 3000 (Thermo Scientific L3000150). Lentiviruses were collected 2 d post-transfection and used to transduce K562 cells. Transduced K562 cells were selected using puromycin at 2 µg ml^-1^. Cell lines were confirmed to express the transduced constructs by cell-surface immunostaining.

### Cell surface staining with Fc fusion proteins and antibodies

HEK293T cells seeded in 6-well plates were transfected with 2 µg expression plasmids encoding wild-type or mutant E3–E2–(6K/TF)–E1 proteins of CHIKV, SFV, or MIDV using Lipofectamine^TM^ 3000 (Thermo Scientific L3000150). 24 h post-transfection, cells were trypsinized, washed in cold wash buffer (PBS with 2% (v/v) goat serum for CHIKV and MIDV; TBS with 2% (v/v) goat serum for SFV) once, and incubated with blocking buffer (PBS with 5% (v/v) goat serum for CHIKV and MIDV; TBS with 5% (v/v) goat serum for SFV) at 4 °C for 30 min. Cells were then washed in cold wash buffer once.

Fc-fusion proteins were diluted in wash buffers to the following concentrations before adding to cells: for cells expressing CHIKV glycoproteins, Lachesin–Fc, *Hs*MXRA8–Fc, and *Mm*MXRA8–Fc were used at 3.16 µg ml^-1^, and anti-SFV immune ascitic fluid (ATCC VR-1247AF) was used at 1:200 dilution; for cells expressing SFV glycoproteins, Lachesin–Fc and VLDLR_LBD_–Fc were used at 1 µg ml^-1^, and anti-SFV immune ascitic fluid was used at 1:200 dilution; for cells expressing MIDV glycoproteins, Lachesin–Fc was used at 100 µg ml^-1^, and humanized macaque antibody SKT05 (hSKT05) (*4*) was used at 10 µg ml^-1^. *Hs*PCDH10_EC1_–Fc was included as a control Fc-fusion protein used at the same concentrations as other Fc-fusion proteins, and an SFV control ascitic fluid (ATCC VR-1247CAF) was included as a control for the immune ascitic fluid used at 1:200 dilution.

Diluted Fc-fusion proteins, hSKT05 antibody, or ascitic fluids were added to cells. Cells were incubated for 30 min at 4 °C. Cells were then washed twice with cold wash buffer. Secondary detection F(ab’)_2_ fragments were diluted at 1:200 in wash buffer: PE-conjugated goat anti-human F(ab’)_2_ fragment (Jackson ImmunoResearch 109-116-098) was used to detect Fc-fusion protein and hSKT05 staining, and PE-conjugated donkey anti-mouse F(ab’)_2_ fragment (Jackson ImmunoResearch 715-116-150) was used to detect ascitic fluid staining. Diluted F(ab’)_2_ fragments were added to cells. Cells were incubated for 30 min at 4 °C. Cells were then washed three times in cold wash buffer, twice in cold PBS for CHIKV and MIDV or cold TBS for SFV, fixed in 2% (v/v) formalin and subjected to detection of surface staining using an iQue3 Screener PLUS (Intellicyt) with ForeCyt (Sartorius) software (version 8.1.7524).

For cell surface immunostaining of K562 cell lines ectopically overexpressing alphavirus receptors, cells were blocked in blocking buffer (5% [v/v] goat serum in PBS) for 30 min at 4 °C and washed once with binding buffer (2% (v/v) goat serum in PBS). Primary antibodies were diluted to 5 µg ml^-1^ in binding buffer. Primary antibodies used include polyclonal anti-VLDLR (GeneTex GTX79552), anti-MXRA8 (MBL International W040-3), anti-Lachesin (*1*), anti-rabbit IgG polyclonal isotype (Proteintech 30000-0-AP), and anti–mouse IgG1 monoclonal isotype (Proteintech 66360-1-Ig). Cells were washed twice in binding buffer and were subsequently incubated with PE-conjugated donkey anti-rabbit F(ab′)2 fragment (Jackson ImmunoResearch 711-116-152) or a PE-conjugated donkey anti-mouse F(ab′)2 fragment (Jackson ImmunoResearch 715-116-150) diluted 1:200 in binding buffer for 30 min at 4 °C. Cells were then washed three times in binding buffer, followed by two washes in PBS before being fixed in 2% (v/v) formalin in PBS. Cell-surface receptor expression was detected using an iQue3 Screener PLUS (Intellicyt) with ForeCyt (Sartorius) software (version 8.1.7524). Antibody staining was visualized using FlowJo (version 10.10.0).

For cell surface immunostaining of K562 cell lines ectopically overexpressing proteins containing a Flag tag, a commercial APC-conjugated anti-Flag (DYKDDDK) antibody (Biolegend 637308) or isotype control (Biolegend 402306) was diluted to 5 µg ml^-1^ in binding buffer as above after a 30 min block in blocking buffer. Cells were then washed three times in binding buffer followed by two washes in PBS before being fixed in 2% (v/v) formalin in PBS. Cell-surface Flag staining was detected using an iQue3 Screener PLUS (Intellicyt) with ForeCyt (Sartorius) software (version 8.1.7524). Antibody staining was visualized using FlowJo (version 10.10.0). For cell surface immunostaining of K562 cells ectopically expressing *Ae. aegypti* LpR1, we incubated cells for one hour with the LDLR family chaperone protein, RAP, containing a C-terminal Flag tag in place of the endoplasmic reticulum retention signal (*27*), diluted to 5 µg ml^-1^ in binding buffer. Following incubation, cells were washed twice in binding buffer before incubation with APC-conjugated anti-Flag (DYKDDDK) antibody (Biolegend 637308) as above. Following anti-Flag antibody incubation, cells were washed three times in binding buffer followed by two washes in PBS before being fixed in 2% (v/v) formalin in PBS. Cell-surface Flag staining was detected using an iQue3 Screener PLUS (Intellicyt) with ForeCyt (Sartorius) software (version 8.1.7524). Antibody staining was visualized using FlowJo (version 10.10.0).

### Reporter virus particle generation

To generate alphavirus RVPs, HEK293T cells were transfected with two plasmids using Lipofectamine 3000 (Thermo Fisher Scientific L3000150) by following the manufacturer’s instructions. One plasmid is a modified pRR64 RRV replicon provided by R. Kuhn (Purdue University) (*57*) in which the SP6 promoter was exchanged with a CMV promoter, and the E3– E2–(6K/TF)–E1 sequence was replaced with a CopGFP (GenBank AAQ01184) reporter preceded by a porcine teschovirus-1 2A self-cleaving peptide. The second plasmid is a pCAGGS or pTWIST-CMV-BG-WPRE-Neo expression vector containing the E3–E2–(6K/TF)–E1 sequence of heterologous alphaviruses. At 4–6 h post-transfection, media was replaced with Opti-MEM supplemented with 5% (v/v) FBS, 25 mM HEPES (Thermo Fisher Scientific), and 5 mM sodium butyrate. Two days after transfection, supernatant was harvested and spun down at 3000 × g for 5 min before being passed through a 0.45 µm filter, aliquoted, and stored at -80 °C. Alphavirus E3–E2–(6K/TF)–E1 sequences included in the pCAGGS vector were CHIKV 37997 (GenBank AY726732), SFV SFV4 (GenBank AKC01668), and MAYV LET-1430 (GenBank PP505832). The alphavirus E3–E2–(6K/TF)–E1 sequences included in the pTWIST-CMV-BG-WPRE-Neo expression vector were MIDV MIDV857 (GenBank EF536323), ONNV SG650 (GenBank YP010775618), RRV T48 (GenBank ACV67002), and GETV M1 (GenBank EU015061).

### Reporter virus particle entry assays

We incubated transduced K562 cells with RVPs. Twenty-four hours post-infection, cells were harvested, washed twice with PBS, and fixed in PBS containing 2% (v/v) formalin. GFP expression was measured by flow cytometry using an iQue3 Screener PLUS (Intellicyt) with IntelliCyt ForeCyt Standard Edition (Sartorius) software (version 8.1.7524). An example of the flow cytometry gating scheme used to quantify GFP-expressing RVP infection is provided in Figure S6C.

For assessment of efficiency of receptor recognition, RVPs were titrated on K562 cells expressing cognate receptors in a twofold dilution series, with MOI calculated based on titers measured on the cell type indicated in the assay description.

### *In vivo* protection study

Cohorts of ten mixed-sex (*n* = 4–5 males and *n* = 5–6 females) six-to eight-week-old A129 mice were obtained from a colony maintained at UTMB. Mice received a 50 mg kg^-1^ dose of either Lachesin–Fc, *Hs*MXRA8–Fc, *Hs*PCDH10_EC1_–Fc, or an equivalent volume of DPBS via the intraperitoneal route. Six hours later, mice were infected with 10^4^ PFU of SFV A774 subcutaneously in the footpad. At two days post-infection, blood was collected via the retro-orbital route and sera were stored at -80°C for subsequent titration via plaque assay. Health checks were performed daily up to 14 days post-infection following SFV challenge.

Cohorts of eight mixed-sex (*n* = 4 males and *n* = 4 females) four-week-old C57BL/6J mice were obtained from Jackson Laboratory (Bar Harbor, ME). Mice received a 50 mg kg^-1^ dose of either Lachesin–Fc, *Hs*MXRA8–Fc, *Hs*PCDH10_EC1_–Fc, or an equivalent volume of DPBS via the intraperitoneal route. Twenty-four hours later, the ipsilateral footpad was measured with a digital caliper immediately before mice were infected with 10^3^ PFU of CHIKV strain SL07 subcutaneously in the footpad. At three days post-infection, the ipsilateral footpad was again measured with a digital caliper, after which the mice were euthanized, and blood was collected immediately prior to transcardial perfusion with DPBS. Contralateral and ipsilateral feet and calf muscle were collected in Vero maintenance media (DMEM with 2% (v/v) FBS, 100 U ml^-1^ penicillin, and 100 µg ml^-1^ streptomycin). All sera and tissue samples were stored at -80 °C for subsequent titration via plaque assay.

All studies were performed in accordance with the NIH Guidance for the Care and Use of Laboratory Animals as approved by the UTMB Institutional Animal Care and Use Committee under protocol 1708051. Mice were monitored at least once daily for the duration of all studies. Sterile water and irradiated 2019 Teklad Global 19% protein extruded rodent diet food pellets (Inotiv, West Lafayette, IN) were provided *ad libitum*. Mice were housed in an environment with 12-hour light cycles, a temperature range of 68–79°F, and a humidity range of 30–70%.

### Plaque assay

Tissue samples were homogenized for two (calf muscle) or three (foot) minutes at 26 s^-1^, and debris was pelleted via centrifugation for five minutes at 16,100 × g prior to titration. Samples were serially one-to-ten diluted in DPBS with 2% (v/v) FBS and allowed to infect monolayers of Vero E6 cells for one hour at 37 °C with 5% CO_2_. After this incubation, an overlay consisting of 0.4% agarose and 0.8× Vero maintenance media was added to the monolayers. Two days later, cells were fixed with 10% neutral buffered formalin and stained with crystal violet. Plaques were counted with the aid of a light box.

### Cryo-EM sample preparation and data collection

Samples were prepared by mixing 3 µl of VLPs (3.0 mg ml^− 1^ in DPBS) with 3 µl of Fc protein (3.0 mg ml^− 1^) and immediately applying 4 µl of the mixture to glow-discharged Quantifoil grids (R 1.2/1.3, 300 mesh, gold; TED PELLA, INC 658-300-AU). Grids were blotted once for 5 s following a 15 s wait at 100% humidity and 4 °C, then plunge-frozen in liquid ethane using a FEI Vitrobot Mark IV (Thermo Fisher Scientific). Cryo-EM data were collected on a 300 kV FEI Titan Krios microscope (Thermo Fisher Scientific) equipped with a Falcon 4 direct electron detector at the Harvard Cryo-Electron Microscopy Center. Automated single-particle acquisition was performed using EPU at 130,000× magnification in counting mode, yielding a calibrated pixel size of 0.94 Å, with datasets collected at defocus values ranging from -0.8 to -2.0 µm.

### Cryo-EM data processing

Raw movie stacks were corrected for beam-induced motion using MotionCor2 (version 1.6.4). (*58*) The parameters of the contrast transfer function (CTF) for all micrographs were estimated by CTFFIND-4.1 (version 4.1.14). (*59*) For the CHIKV VLP in complex with Lachesin–Fc, 14,402 particles were auto-picked from 13,138 micrographs using crYOLO (version 1.8.2) (*60*) and particle extraction was carried out in RELION (version 3.1) (*61*). Several rounds of reference-free 2D classification yielded 10,758 high-quality particles for 3D classification with icosahedral symmetry, using the 12 Å density map of WEEV VLP (EMD-5210) (*62*) as the initial reference. A subset of 5,941 particles was selected for 3D auto-refinement with icosahedral symmetry, resulting in a 6.4 Å reconstruction of the CHIKV VLP:Lachesin–Fc complex.

To further improve map resolution, a block-based reconstruction approach (*30*) centered on the q3 E2–E1 trimer was applied. In this step, 356,460 particle blocks, each containing four E2–E1 trimers near quasi-threefold axes (q3), were extracted and classified without alignment. Following 3D classification, 177,416 selected blocks underwent 3D auto-refinement with a soft mask around the asymmetric unit, producing a density map at 3.4 Å. The final map was post-processed using DeepEMhancer (version 20241203) (*63*) for model building. Details related to workflow, image numbers, and particle counts are presented in Figure S1 and Table S1.

For CHIKV VLP:HsMXRA8–Fc, SFV VLP:Lachesin–Fc, and MIDV VLP:Lachesin–Fc complexes, similar data processing steps were performed. Resolution estimations used the gold standard Fourier shell correlation (FSC) at the 0.143 criterion, with local resolution of the density maps of trimeric virus-receptor complex determined using Relion (*61*).

### Model building and refinement

To build the structural model of the CHIKV VLP:Lachesin–Fc complex, existing structures of CHIKV proteins (E1, E2, and capsid protein; PDB: 6NK5) and the Lachesin structure predicted by AlphaFold (*3*) were used as references. These models were initially docked into the 3.4 Å cryo-EM density map using UCSF ChimeraX (version 1.9) (*64*) then manually rebuilt and adjusted in Coot (version 0.9.8.91) (*65*) followed by iterative real-space refinements in Phenix (version 1.21rc1-5127) (*66*). For the CHIKV VLP:*Hs*MXRA8–Fc complex, CHIKV structural proteins (PDB: 6NK5) (*25*) and the HsMXRA8 structure (PDB: 6JO8) (*43*) were used as starting models. Similarly, for the SFV VLP:Lachesin–Fc complex, structures of SFV proteins (PDB: 8UA8) (*5*) and the Lachesin model from the CHIKV complex were used. For the MIDV VLP:Lachesin–Fc complex, AlphaFold-predicted MIDV proteins and the Lachesin model from CHIKV complex served as references. All models were built and refined using Coot (version 0.9.8.91) (*65*) and Phenix (version 1.21rc1-5127) (*66*). For the structure of CHIKV bound to human MXRA8–Fc, coordinates resembled those previously reported for CHIKV bound to MXRA8 structures, with Cα RMSDs of 0.70 Å and 0.90 Å relative to the cryo-EM structure (PDB ID: 6NK6) (*25*) and the X-ray crystal structure (PDB ID: 6JO8) (*43*), respectively. Data collection, refinement, and validation statistics are summarized in Table S1. Structural analyses and figure preparation were conducted using ChimeraX (version 1.9) (*64*) and PyMOL (version 3.1.8, https://www.pymol.org/pymol). Software tools were curated by SBGrid. (*67*)

### Sequence alignment of Lachesin orthologs

Sequence alignment of Lachesin orthologs of *Aedes albopictus* (GenBank: XP_019932516), *Aedes aegypti* (GenBank: XP_001659911), *Wyeomyia smithii* (GenBank: XP_055534723), *Uranotaenia lowii* (GenBank: XP_055609662), *Culex quinquefasciatus* (GenBank: XP_001850358), *Culex pipiens pallens* (GenBank: XP_039444183), *Anopheles darlingi* (GenBank: XP_049542368), *Anopheles coustanni* (GenBank: XP_058121093), *Phlebotomus papatasi* (GenBank: XP_055707817), *Apis mellifera* (GenBank: XP_397471), *Ixodes scapularis* (GenBank: XP_029832537) was performed using the MAFFT program in the EMBL-EBI Job Dispatcher (*68*) and visualized using ESpript3.0 (*69*).

### Phylogenetic tree generation

For the tree of alphaviruses, amino acid sequences of the E2 proteins of the following alphaviruses were aligned in MEGA12 (version 12.0.14) using the built-in MUSCLE algorithm: CHIKV 37997 (GenBank AAU43881), SFV SFV4 (GenBank AKC01668), RRV T48 (GenBank ACV67002), GETV M1 (GenBank EU015061), MAYV LET-1430 (GenBank PP505832), UNAV BeAr13136 (GenBank AF339481), BEBV MM2354 (GenBank AF339480), ONNV SG650 (GenBank YP010775618), MIDV MIDV857 (GenBank EF536323), BFV K61404 (GenBank MN689027), WEEV 71V1658 (GenBank GQ287645), VEEV INH-9813 (GenBank KP282671), and EEEV FL91-469 (GenBank AY705241). A maximum-likelihood phylogenetic tree was constructed for the aligned sequences using the LG model with discrete Gamma distribution across four categories. The bootstrap method was used to test phylogeny with 1000 bootstrap replicates.

**Figure S1.**
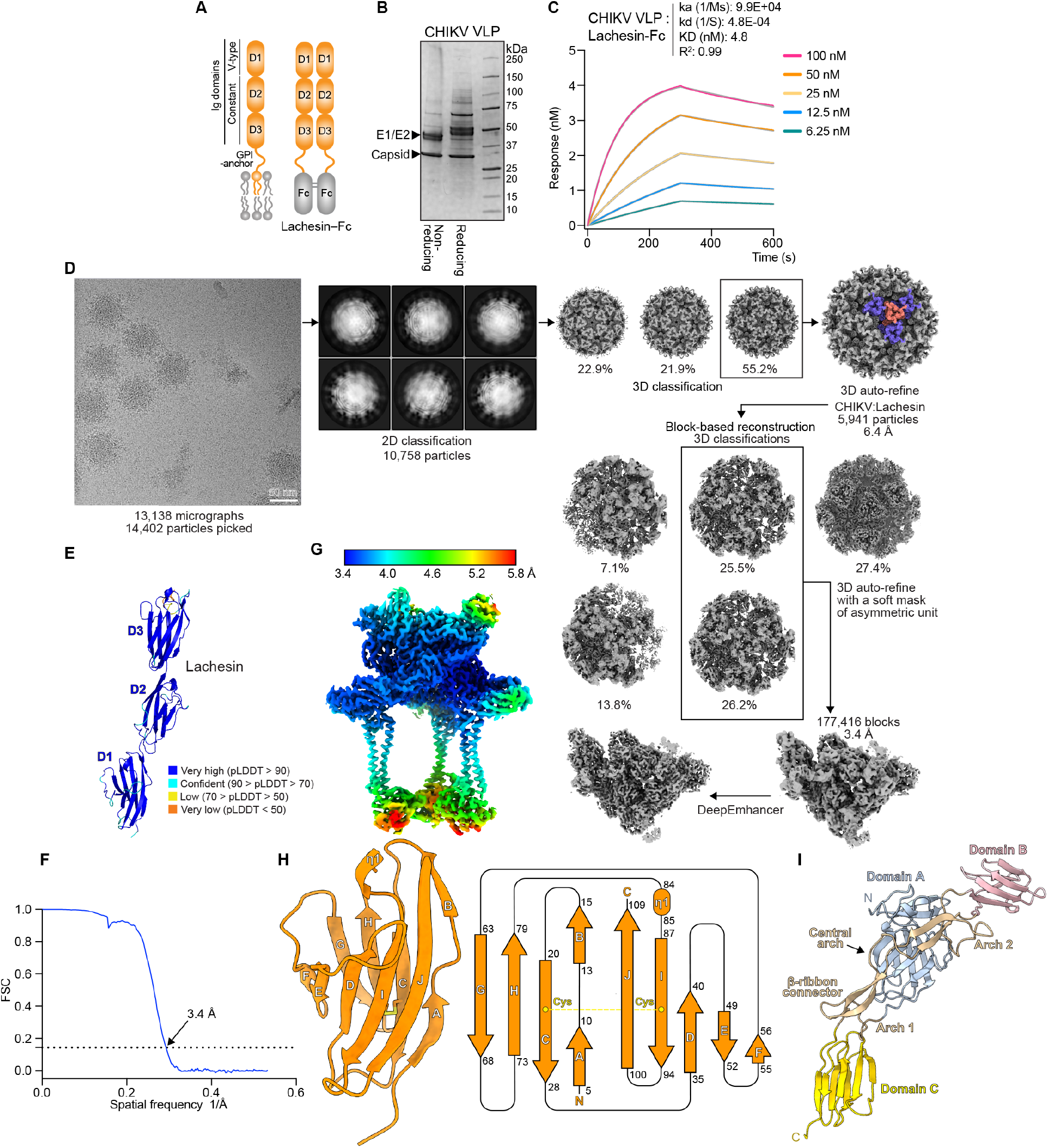
Cryo-EM data processing of CHIKV VLPs in complex with Lachesin–Fc. **(A)** Schematic diagrams of Lachesin and Lachesin–Fc construct. **(B)** Coomassie-stained SDS-PAGE gel of purified CHIKV 37997 VLPs. **(C)** Biolayer interferometry binding analysis of Lachesin–Fc with immobilized CHIKV VLPs. Data are colored as indicated for the different concentrations and the fit is shown in gray. Kinetic parameters of binding were determined using a 1:1 interaction model. **(D)** Workflow used for cryo-EM data processing of CHIKV VLPs bound to Lachesin–Fc. **(E)** Model of Lachesin D1–D3 predicted by AlphaFold (*3*) shown in ribbon representation and colored by Predicted Local Distance Difference Test (pLDDT) score ranges. **(F)** Fourier shell correlation curves of CHIKV VLP in complex with Lachesin–Fc. The threshold used to estimate the resolution is 0.143. **(G)** Local resolution map of CHIKV with Lachesin–Fc complex estimated using Relion (*61*). **(H)** Ribbon and topology diagram showing the structure of Lachesin D1 with β-strands labeled using the standard Ig-like fold convention. The β-strands are depicted as a cartoon with residue number start/stop locations. Disulfide bonds between cysteines are indicated. **(I)** Schematic diagram of CHIKV E2. The central arch of the β-ribbon connector is denoted. Domains are as established by Voss *et al*. (*70*).

**Figure S2.**
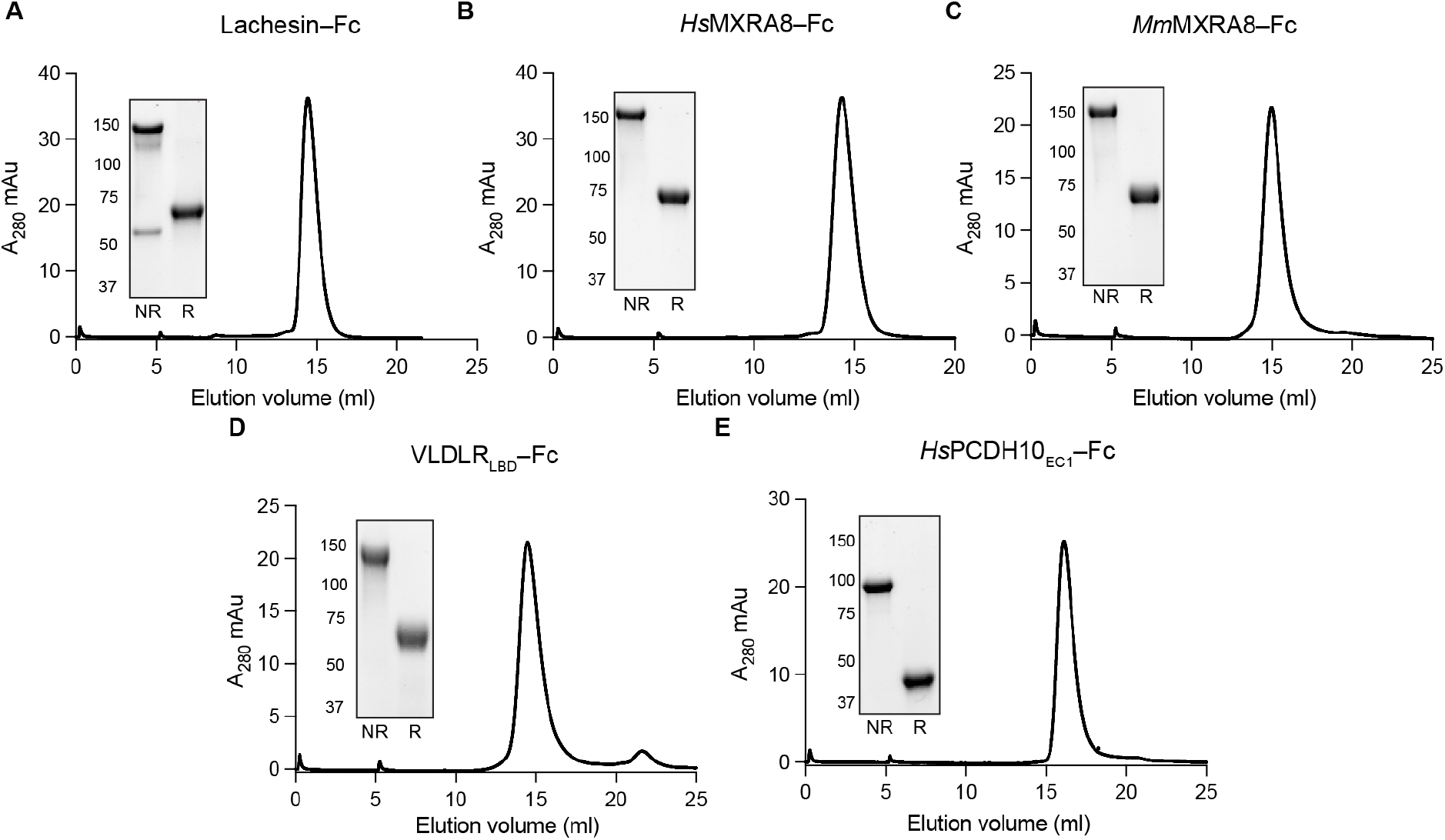
Size-exclusion chromatography traces for Fc fusion proteins and cell line immunostaining experiments. **(A–E)** Size-exclusion chromatography (SEC) trace of Lachesin–Fc (**A**), *Hs*MXRA8–Fc (**B**), *Mm*MXRA8–Fc (**C**), human VLDLR_LBD_–Fc (**D**), or *Hs*PCDH10EC1–Fc (**E**). Insets are Coomassie-stained SDS-PAGE gels.

**Figure S3.**
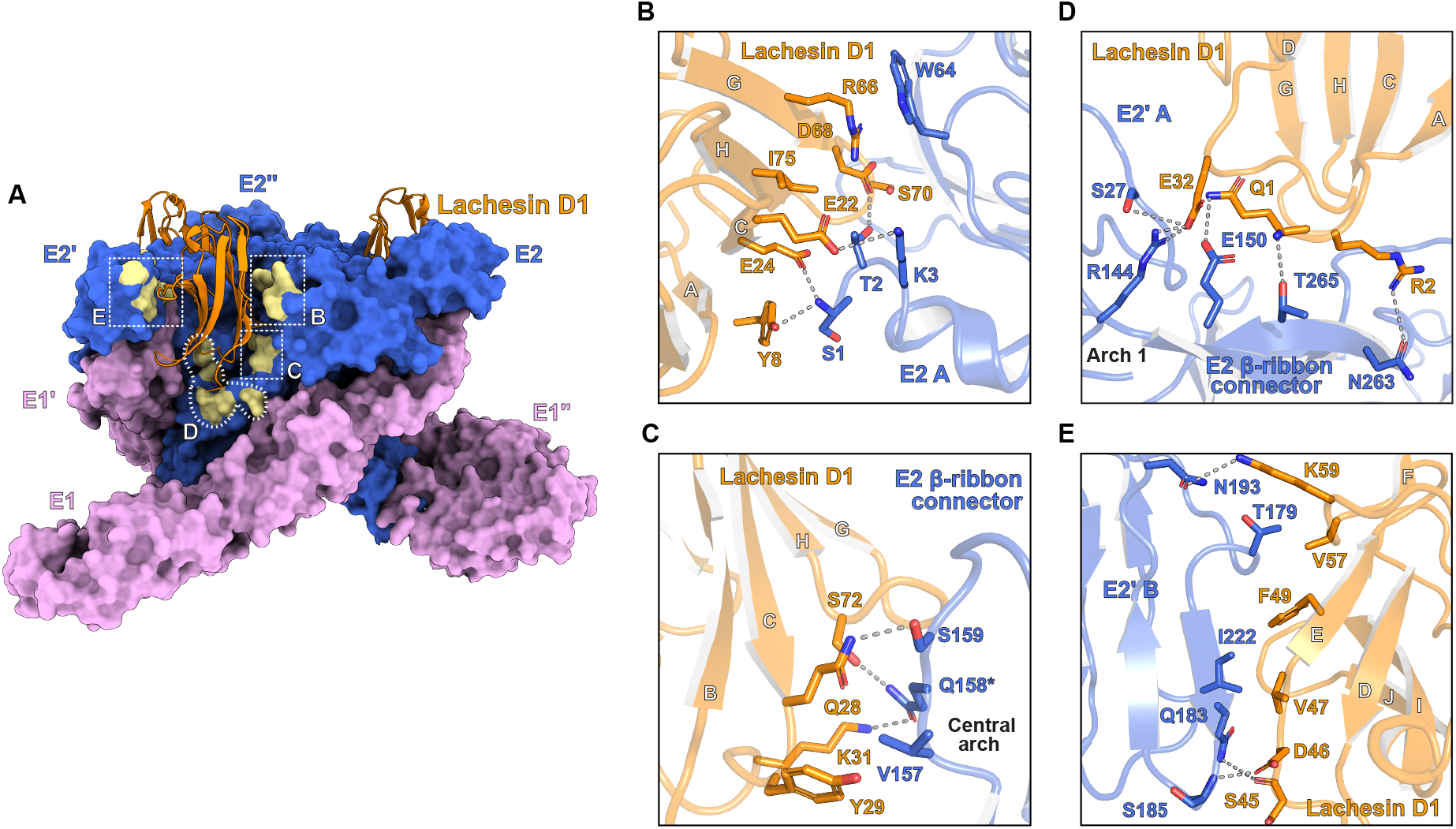
Detailed *Ae. albopictus* Lachesin interactions with CHIKV E2–E1. **(A)** Cryo-EM structure of CHIKV trimer bound to *Ae. albopictus* Lachesin D1. The E2–E1 glycoproteins are shown in surface representation, and the receptor is shown as a ribbon diagram. Annotation is provided for the zoom-in views shown in panels B–E. **(B–E)**, Zoom-in view of regions highlighting Lachesin D1 interactions with a CHIKV E2–E1 heterodimer and its adjacent E2’–E1’ heterodimer. Polar contacts are shown as dashed lines. Contacts include those made by Lachesin R66 in βG, which makes cation-π stacking interactions with E2 domain A residue W64 (**B**). Lachesin residues Q28, Y29, and K31 in loop C–D and S72 in loop G–H participate in polar and hydrophobic interactions with E2 residues V157, Q158, and S159, which are found in the central arch of the β-ribbon connector (**C**). N-terminal Lachesin residues Q1 and R2 are part of an extensive network of polar interactions with the E2 β-ribbon connector (**D**), and Lachesin also makes several polar and hydrophobic contacts with domains A and B of the adjacent E2’ protomer (**D** and **E**). Polar contacts are shown as dashed lines.

**Figure S4.**
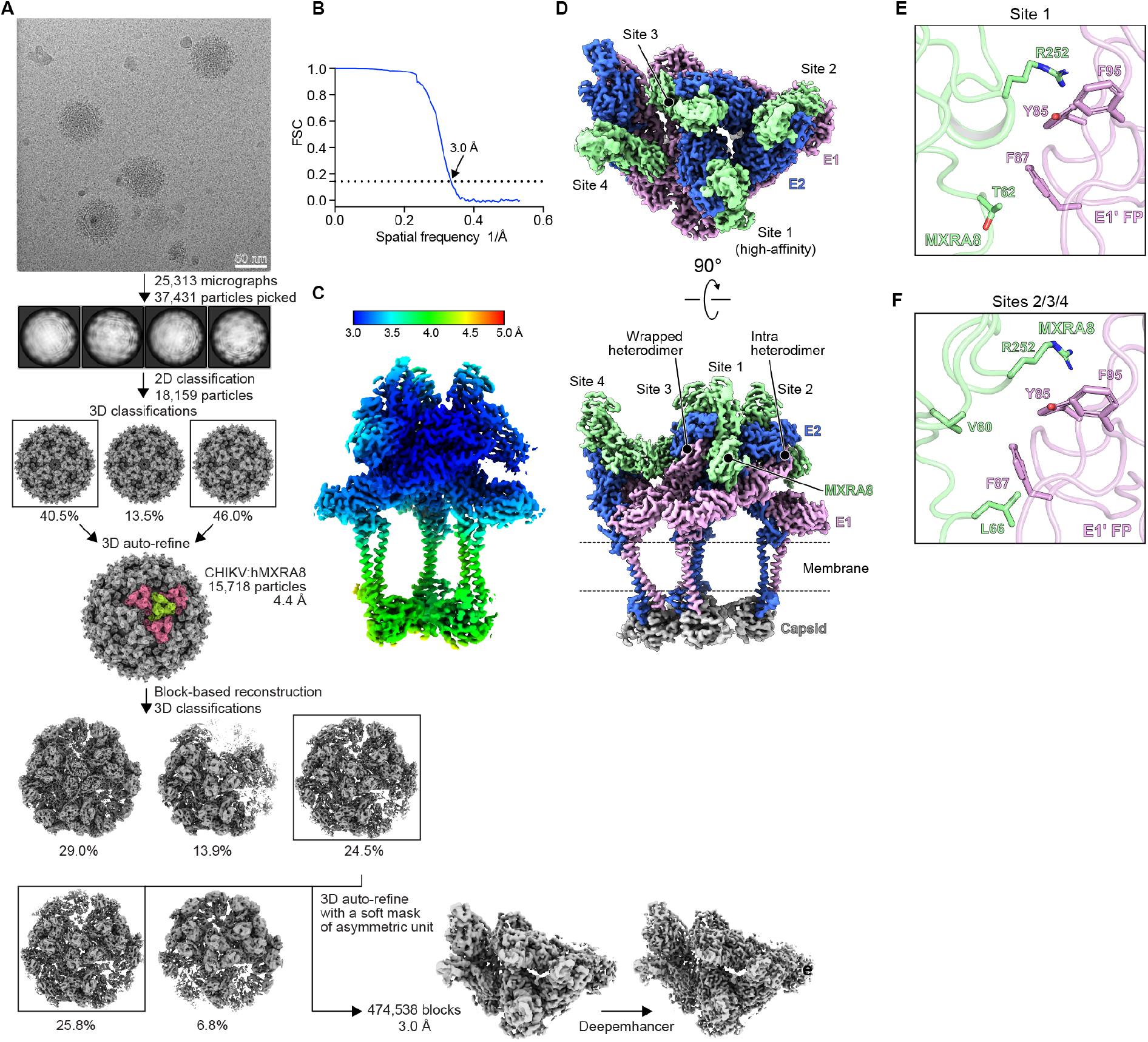
Cryo-EM reconstruction of CHIKV VLP in complex with *Hs*MXRA8–Fc. **(A)** Workflow used for cryo-EM data processing of CHIKV 37997 VLPs bound to *Hs*MXRA8–Fc. **(B)** Fourier shell correlation curves of CHIKV VLP bound to *Hs*MXRA8–Fc. The threshold used to estimate the resolution is 0.143. **(C)** Local resolution map of CHIKV with *Hs*MXRA8–Fc complex estimated using Relion (*61*). **(D)** Focused reconstruction of the asymmetric unit of CHIKV VLP in complex with human MXRA8, colored as in Figure 2A. The location of the high-affinity binding site (site 1) as described by Basore et al. (*25*) and the other sites are indicated. In the high-affinity binding mode (site 1), MXRA8 makes 22 E2–E1 contacts, while in the low-affinity binding mode (sites 2–4), MXRA8 makes 20 contacts. See also Figure S5. **(E**,**F)** Residues of MXRA8 contacting the fusion loop of CHIKV 37997 VLP E1 in site 1 (**E**) and sites 2–4 (**F**).

**Figure S5.**
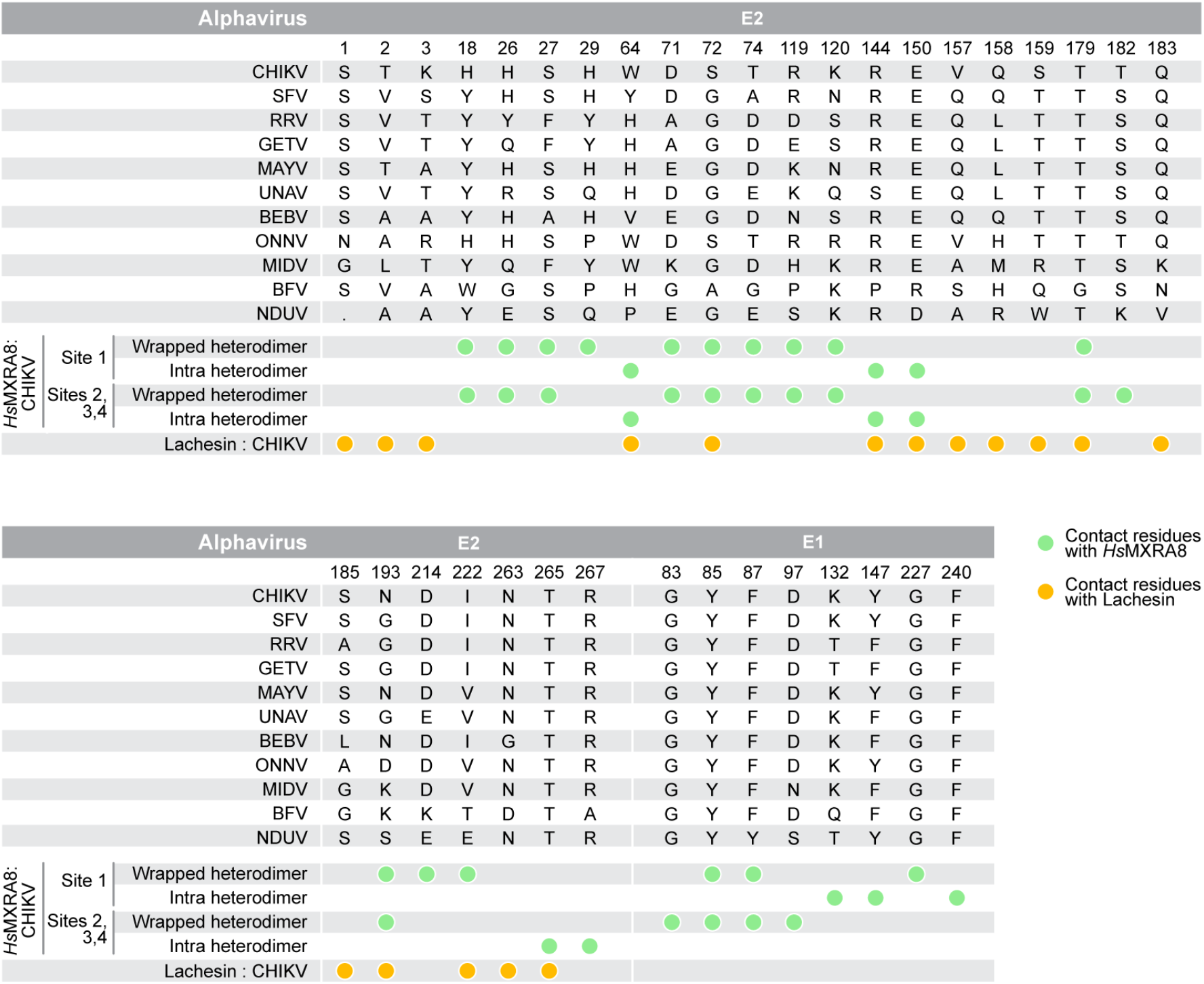
List of residues on CHIKV that contact Lachesin and MXRA8. **(A)** Contact residues between CHIKV E2–E1 and *Ae. albopictus* Lachesin or human MXRA8 (*Hs*MXRA8) from the cryo-EM structures reported in the manuscript are indicated. The “intra” and “wrapped” heterodimer naming convention is from Basore et al. (*25*).

**Figure S6.**
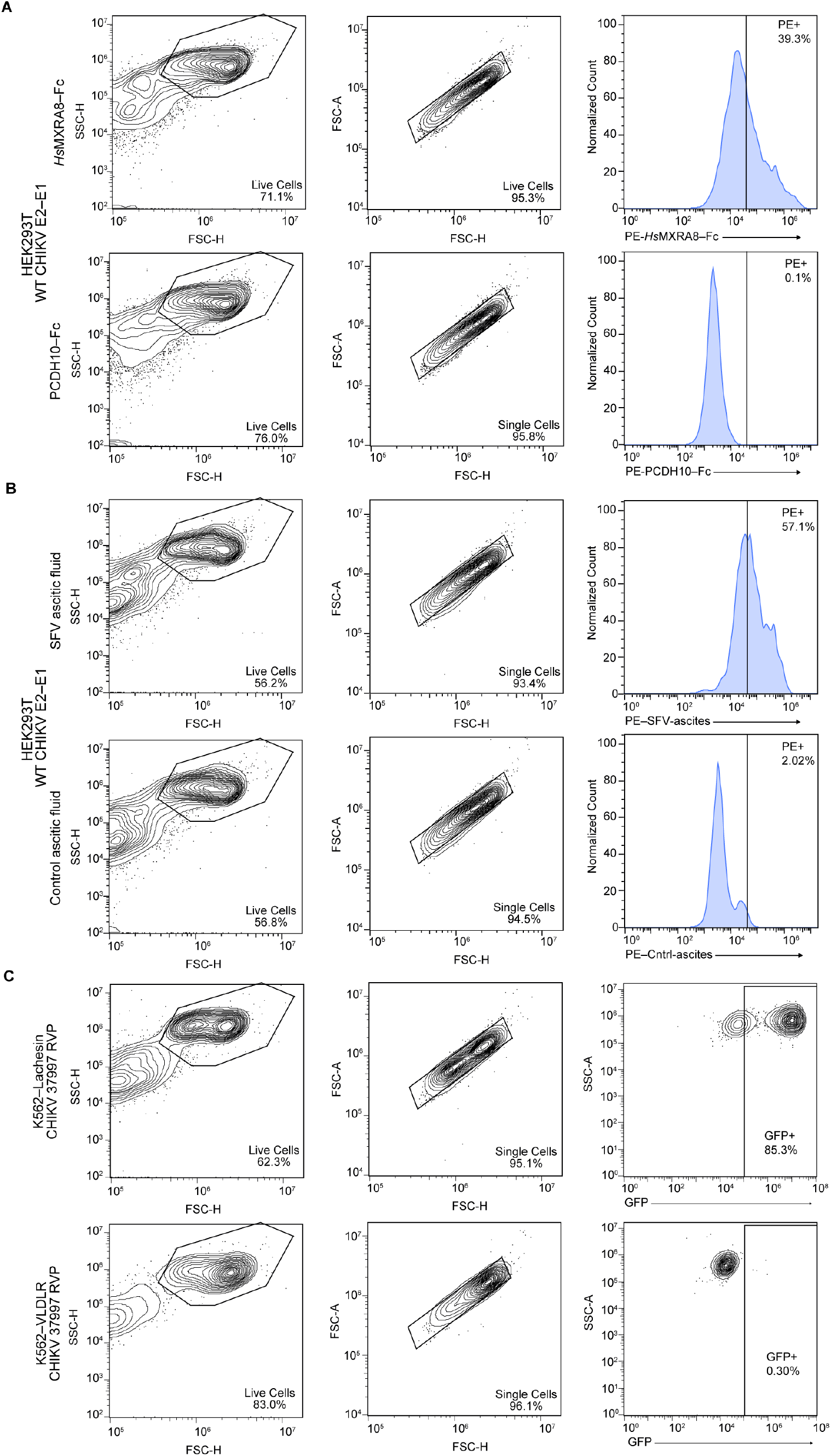
Cell surface staining and GFP expression gating strategies. **(A)** Gating strategy for cell surface staining of HEK293T cells transfected with alphavirus E2–E1 glycoproteins with Fc fusion proteins. HEK293T cells transfected with WT CHIKV 37997 E2–E1 were stained with *Hs*MXRA8– Fc (top) or *Hs*PCDH10_EC1_–Fc (bottom) and detected with a PE conjugated secondary antibody. PE: *R*-phycoerythrin. **(B)** Gating strategy for cell surface staining of HEK293T cells transfected with alphavirus E2–E1 glycoproteins with ascitic fluid controls for expression normalization. HEK293T cells transfected with WT CHIKV 37997 E2–E1 were stained with SFV ascitic fluid (top) or Control ascitic fluid (bottom) and detected with a PE conjugated secondary antibody. PE: *R*-phycoerythrin. **(C)** Gating strategy for RVP-infected K562 cells. K562 cells ectopically overexpressing *Ae. albopictus* Lachesin (top) or VLDLR (bottom) infected with CHIKV 37997 RVPs at an MOI of 2 as titered on K562–Lachesin cells.

**Figure S7.**
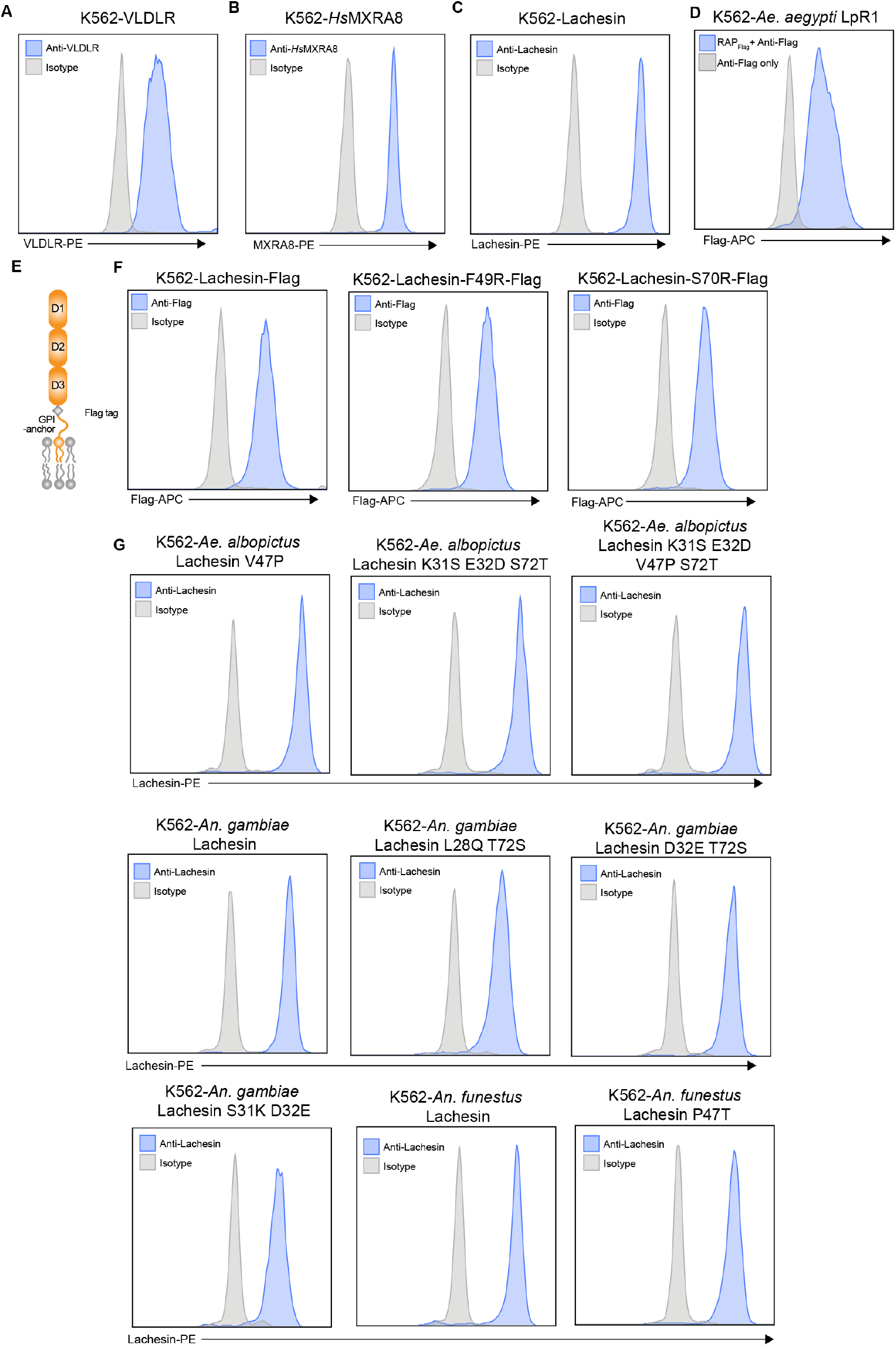
Cell surface immunostaining of K562 cells expressing wild-type and mutant receptors. **(A–D)** Immunostaining of K562 cells stably expressing human VLDLR (**A**), human MXRA8 (**B**), *Ae. albopictus* Lachesin (**C**), or *Ae. aegypti* LpR1 (**D**). **(E)** Schematic diagram of the Lachesin–Flag construct, indicating the position of the internal Flag tag. **(F)** Immunostaining of K562 cells expressing Lachesin–Flag or Lachesin–Flag containing the F49R or S70R substitutions. **(G)** Immunostaining of K562 cells expressing the indicated wild-type or mutant mosquito orthologs of Lachesin.

**Figure S8.**
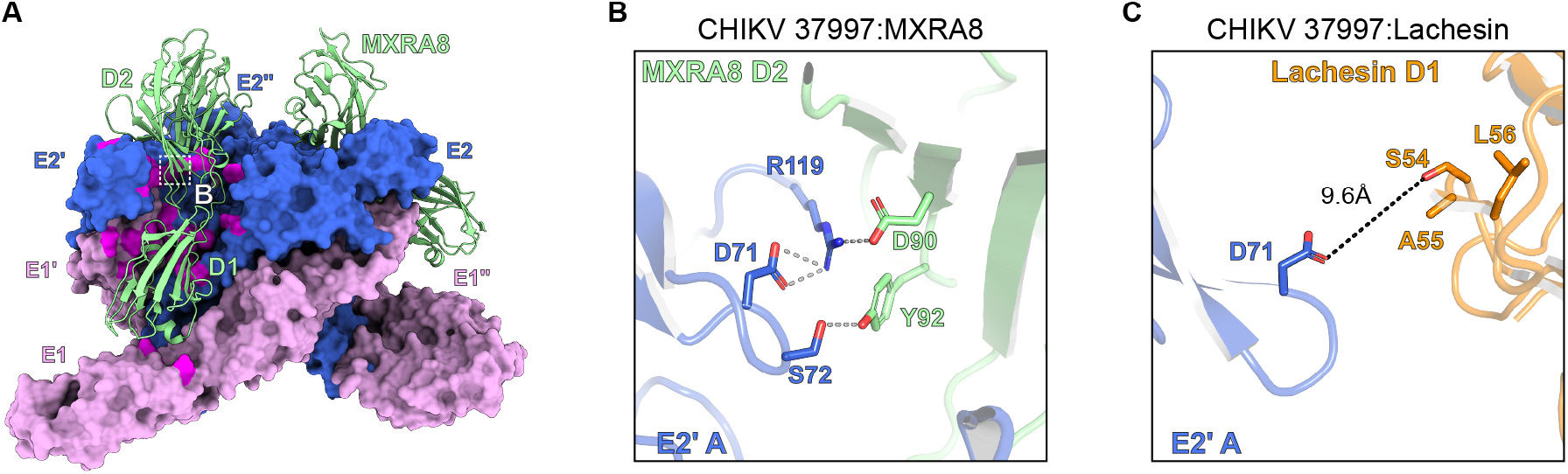
CHIKV E2 residue D71 interacts with MXRA8 but not with Lachesin. **(A)** Cryo-EM structure of CHIKV trimer bound to human MXRA8. The E2–E1 glycoproteins are shown in surface representation, and the receptor is shown as a ribbon diagram. Annotation is provided for the zoom-in view shown in panel B. **(B)** Zoom-in view for the region from the box in (**A**) shown for the CHIKV E2′ domain A residue D71 interaction with neighboring residues and indirect interactions with human MXRA8. Polar contacts are shown as dashed lines. **(C)** The corresponding region to **B** in the CHIKV E2–E1 Lachesin-bound structure is shown. The distance between D71 and the closest Lachesin residue is shown.

**Figure S9.**
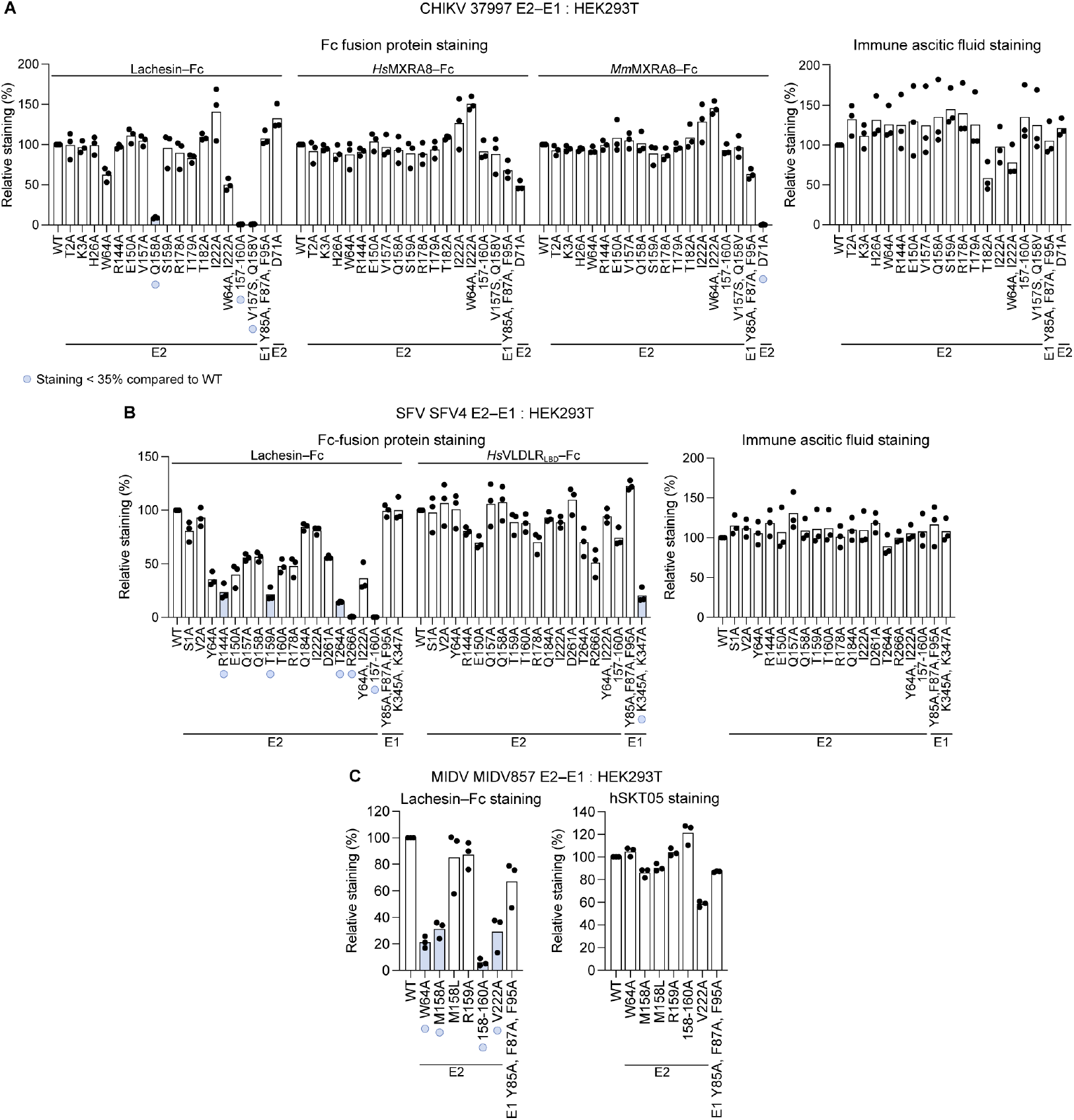
Cell surface immunostaining of E2–E1 mutants with Fc fusion proteins. **(A)** HEK293T cell-surface staining of mutant CHIKV E2–E1 glycoproteins by Lachesin–Fc, *Hs*MXRA8–Fc, or *Mm*MXRA8–Fc fusion proteins normalized to staining levels of WT CHIKV E2–E1 glycoproteins (left) and HEK293T cell-surface staining of mutant CHIKV E2–E1 glycoproteins by anti-SFV immune ascitic fluid (right). Mutants with relative staining <35% of WT levels are highlighted in light blue and indicated with light blue dots. Data are mean from three independent experiments. **(B)** HEK293T cell-surface staining of mutant SFV E2–E1 glycoproteins by Lachesin–Fc or VLDLR_LBD_–Fc fusion proteins normalized to staining levels of WT SFV E2–E1 glycoproteins (left) and HEK293T cell-surface staining of mutant SFV E2–E1 glycoproteins by anti-SFV immune ascitic fluid (right). Mutants with relative staining <35% of WT levels are highlighted in light blue and indicated with light blue dots. Data are mean from three independent experiments. **(C)** HEK293T cell-surface staining of mutant MIDV E2–E1 glycoproteins by Lachesin–Fc normalized to staining levels of WT MIDV E2– E1 glycoproteins (left) and HEK293T cell-surface staining of mutant MIDV E2–E1 glycoproteins by hSKT05, a humanized version of a macaque antibody that cross-reacts with the E1 glycoproteins of alphaviruses (*4*) (right). Mutants with relative staining <35% of WT levels are highlighted in light blue and indicated with light blue dots. Data are mean from three independent experiments.

**Figure S10.**
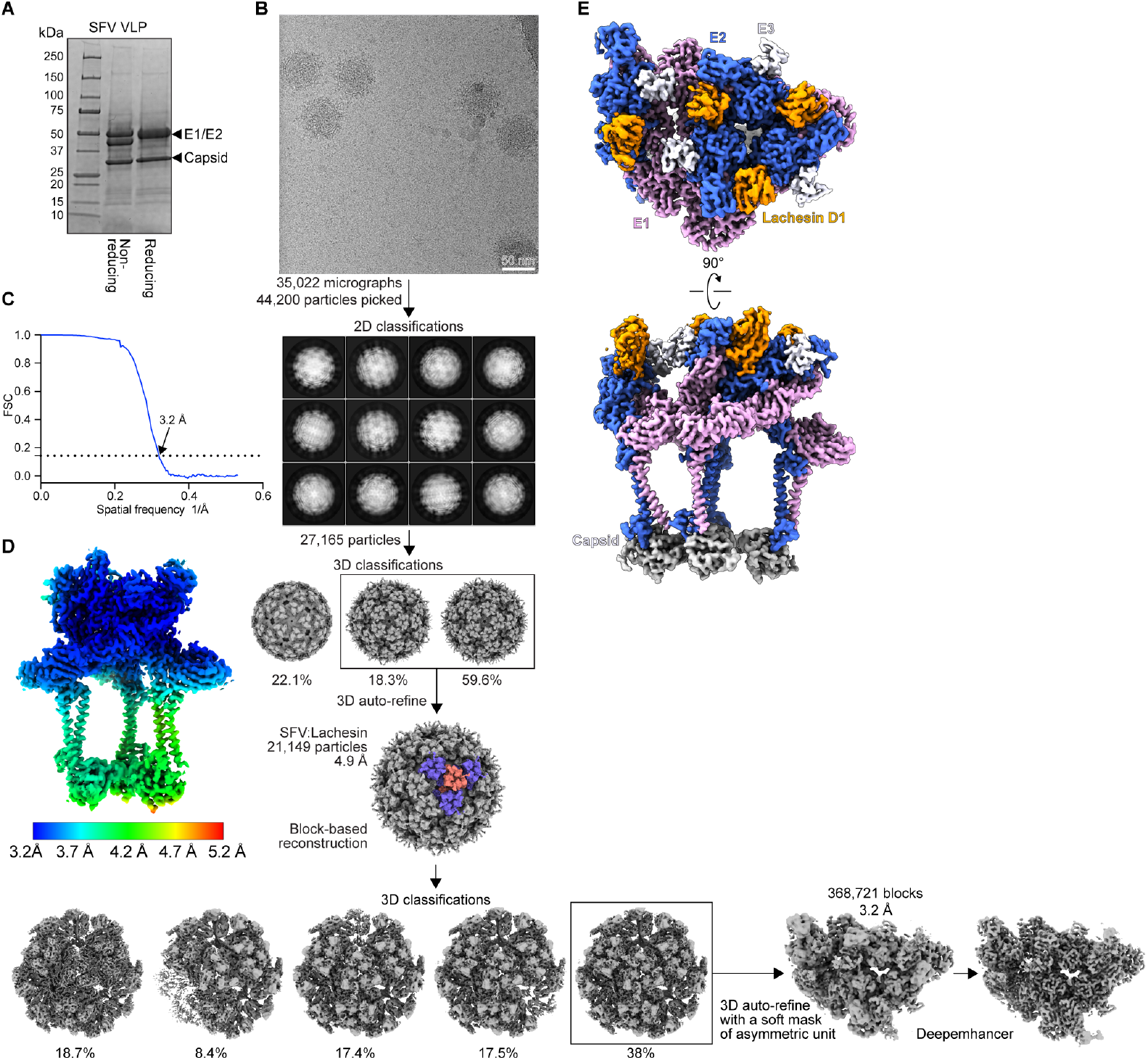
Cryo-EM reconstructions of SFV VLP in complex with Lachesin–Fc. **(A)** Coomassie-stained SDS-PAGE gel of purified SFV VLPs. **(B)** Workflow used for cryo-EM data processing of SFV VLPs bound to Lachesin–Fc. **(C)** Fourier shell correlation curves of SFV VLP bound to Lachesin–Fc. The threshold used to estimate the resolution is 0.143. **(D)** Local resolution map of SFV with Lachesin–Fc complex estimated using Relion (*61*). **(E)** Focused reconstruction of SFV asymmetric unit (ASU) in complex with Lachesin in top view (top) and side view (bottom).

**Figure S11.**
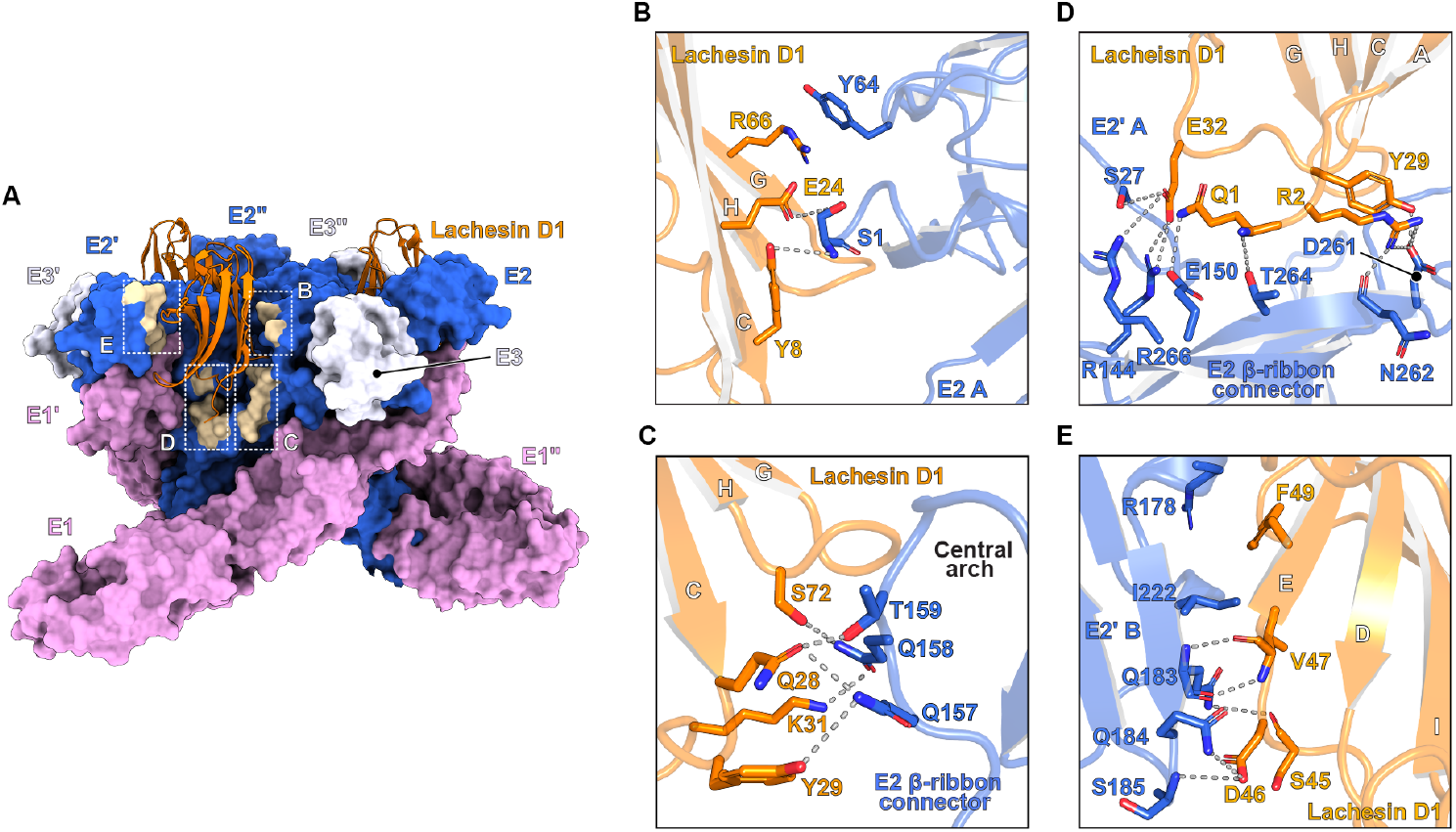
Detailed *Ae. albopictus* Lachesin interactions with SFV E2–E1. **(A)** SFV E2–E1 trimer bound to Lachesin D1. The SFV E2–E1 trimers and their associated E3 proteins are surface rendered, and Lachesin D1 molecules are shown as ribbon diagrams. SFV E2–E1 residues that interact with D1 are in beige. **(B–E)** Zoomed-in view of regions that are boxed in panel **C**. Polar contacts are shown as dashed lines. Interface contacts are reorganized when the Lachesin-bound CHIKV and SFV structures are compared. For example, in the CHIKV-receptor-bound structure, β-ribbon connector residue V157 makes hydrophobic contacts with Lachesin Y29, whereas in SFV, the corresponding E2 residue Q157 makes polar contacts with Lachesin residues Q28 and Y29 (**B**). Additionally, while CHIKV E2′ domain B interactions with Lachesin are dominated by hydrophobic contacts, SFV E2′ domain B interactions with Lachesin are dominated by mostly polar contacts (**E**).

**Figure S12.**
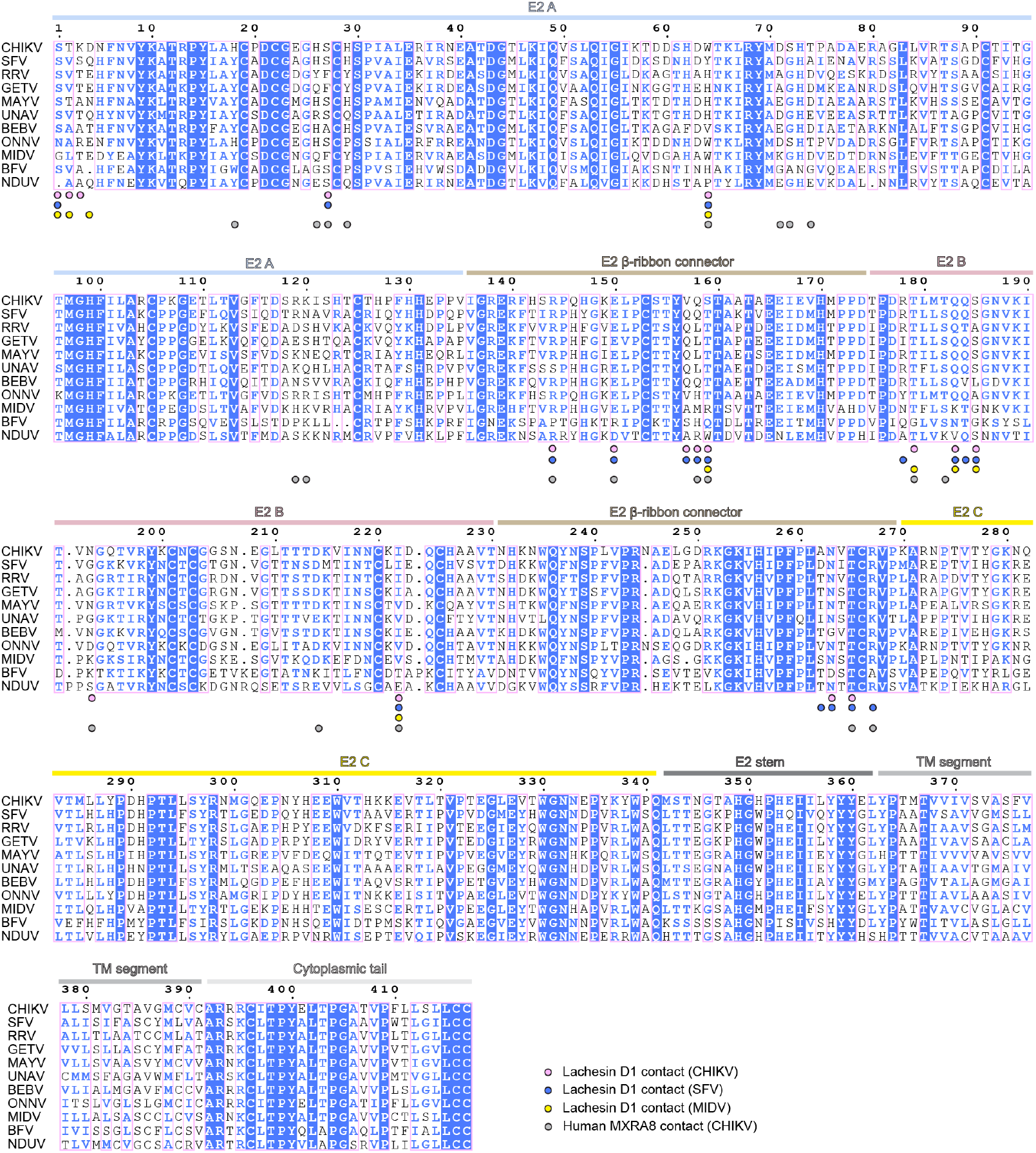
Sequence alignment of alphavirus E2 glycoproteins. The strain information and accession numbers are as follows: CHIKV 37997 (GenBank AAU43881), SFV SFV4 (GenBank AKC01668), RRV T48 (GenBank ACV67002), GETV M1 (GenBank EU015061), MAYV LET-1430 (GenBank PP505832), UNAV BeAr13136 (GenBank AF339481), BEBV MM2354 (GenBank AF339480), ONNV SG650 (GenBank YP010775618), MIDV MIDV857 (GenBank EF536323), NUDV SaAr2204 (GenBank QQZ00849), and BFV K61404 (GenBank MN689027). The blue background denotes residues that are completely conserved in all sequences. Boxed residues indicate regions where a single majority residue or multiple chemically similar residues are found, and these residues are shown in blue font. All other residues are shown in black font.

**Figure S13.**
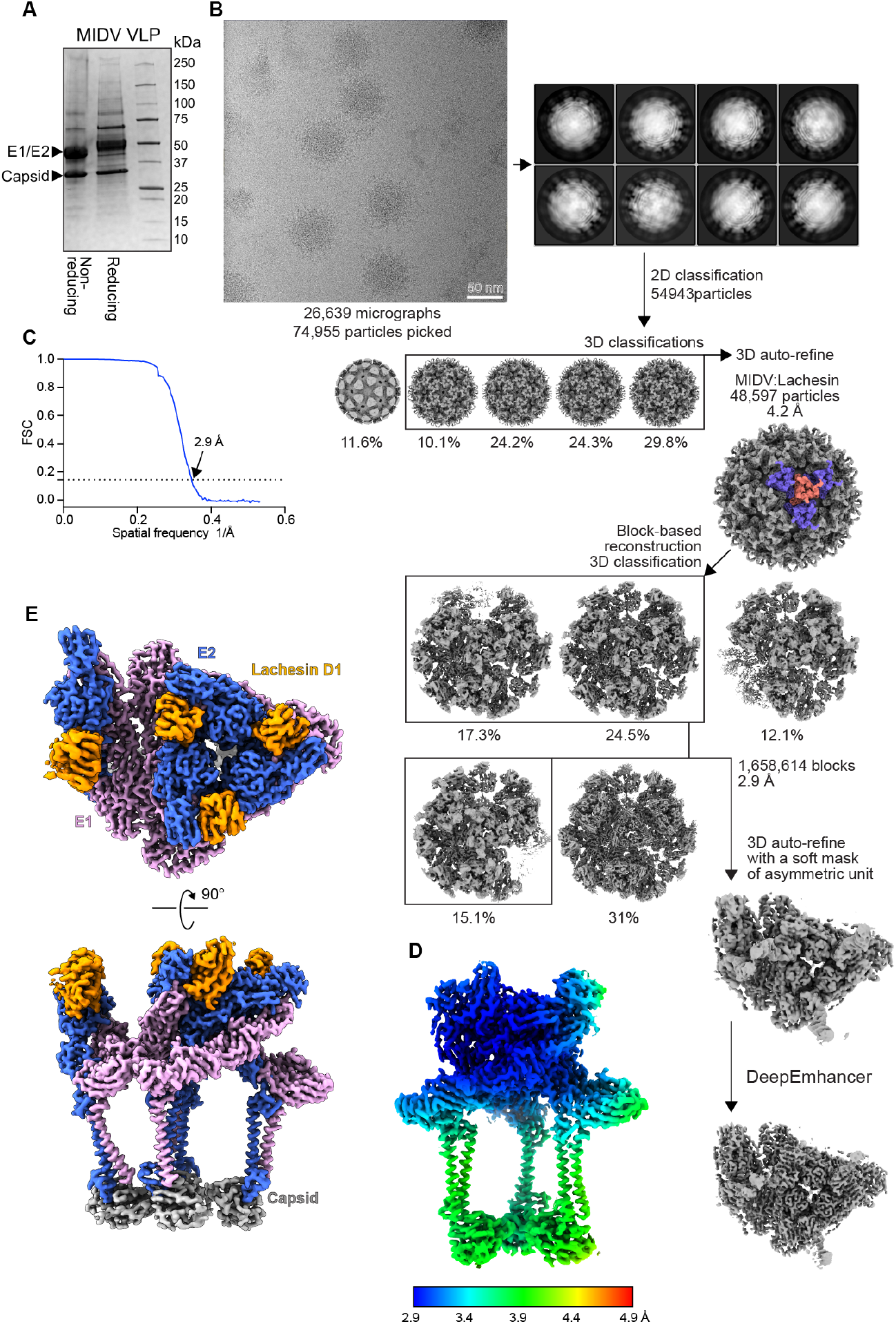
Cryo-EM data processing of MIDV VLPs bound to Lachesin–Fc. **(A)** Coomassie-stained SDS-PAGE gel of purified MIDV VLPs. **(B)** Workflow used for cryo-EM data processing of MIDV VLPs bound to Lachesin–Fc. **(C)** Fourier shell correlation curves of MIDV VLP bound to Lachesin–Fc. The threshold used to estimate the resolution is 0.143. **(D)** Local resolution map of MIDV with Lachesin–Fc complex estimated using Relion (*61*). **(E)** Focused reconstruction of MIDV ASU in complex with Lachesin in top view (top) and side view (bottom).

**Figure S14.**
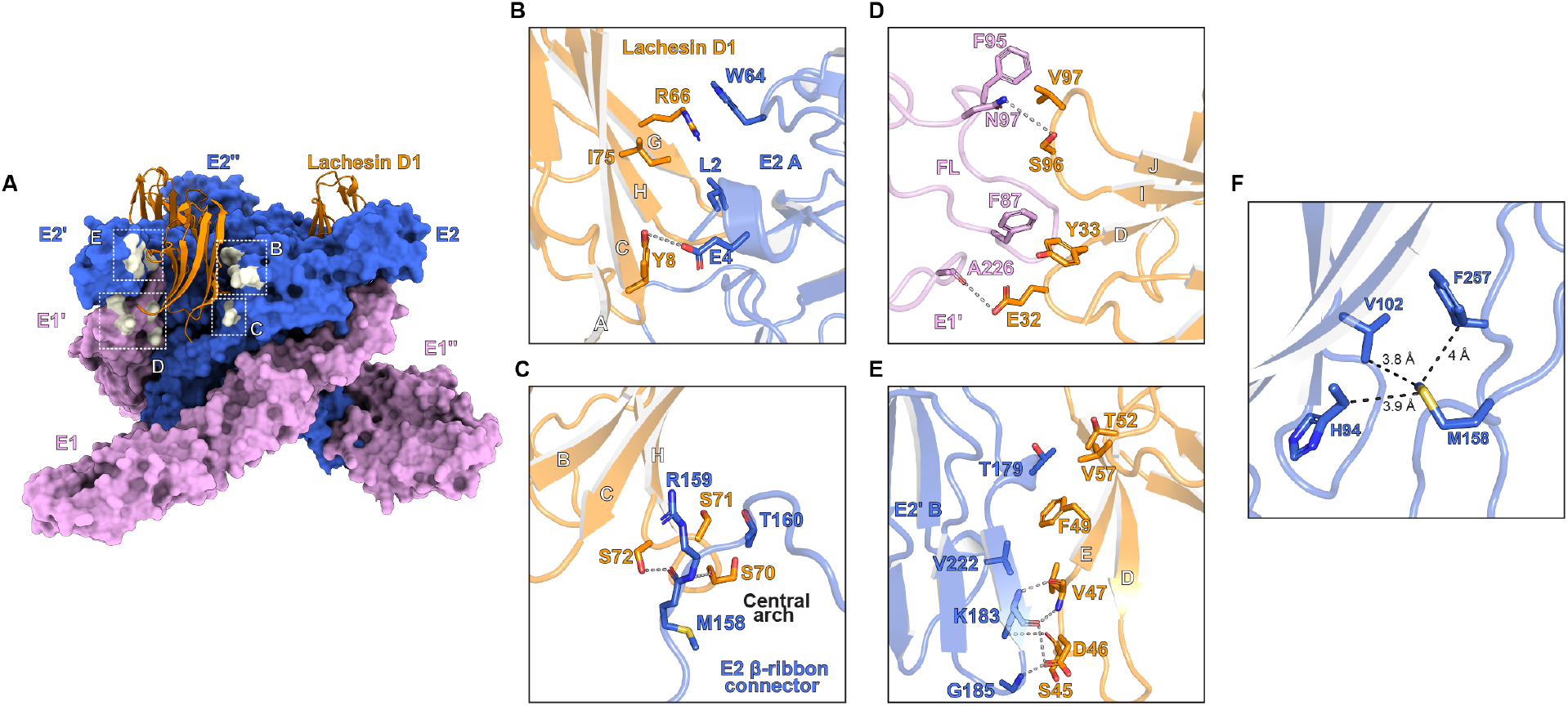
Detailed *Ae. albopictus* Lachesin interactions with MIDV E2–E1. **(A)** MIDV E2–E1 trimer bound to Lachesin D1. MIDV E2–E1 residues that interact with D1 are shown in beige. **(B–E)** Zoom-in view of regions from boxes shown in A. Polar contacts are shown as dashed lines. **(F)** Hydrophobic interactions between MIDV E2 M158 and nearby residues on MIDV E2. Distances between the M158 side chain and surrounding hydrophobic residues are indicated.

**Figure S15.**
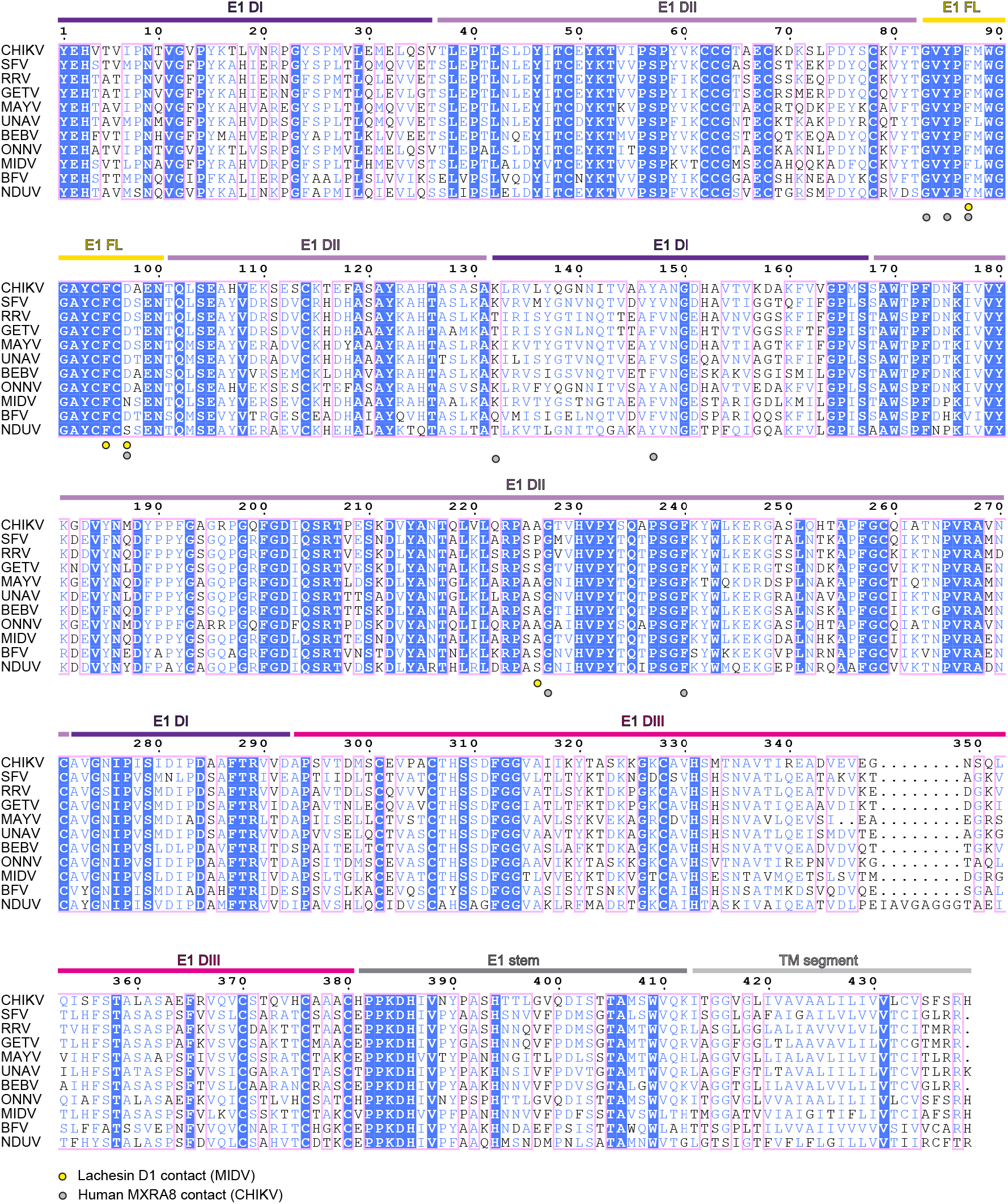
Sequence alignment of alphavirus E1 glycoproteins. The strain information and accession numbers are as follows: CHIKV 37997 (GenBank AAU43881), SFV SFV4 (GenBank AKC01668), RRV T48 (GenBank ACV67002), GETV M1 (GenBank EU015061), MAYV LET-1430 (GenBank PP505832), UNAV BeAr13136 (GenBank AF339481), BEBV MM2354 (GenBank AF339480), ONNV SG650 (GenBank YP010775618), MIDV MIDV857 (GenBank EF536323), NUDV SaAr2204 (GenBank QQZ00849), and BFV K61404 (GenBank MN689027). The blue background denotes residues that are completely conserved in all sequences. Boxed residues indicate regions where a single majority residue or multiple chemically similar residues are found, and these residues are shown in blue font. All other residues are shown in black font.

**Figure S16.**
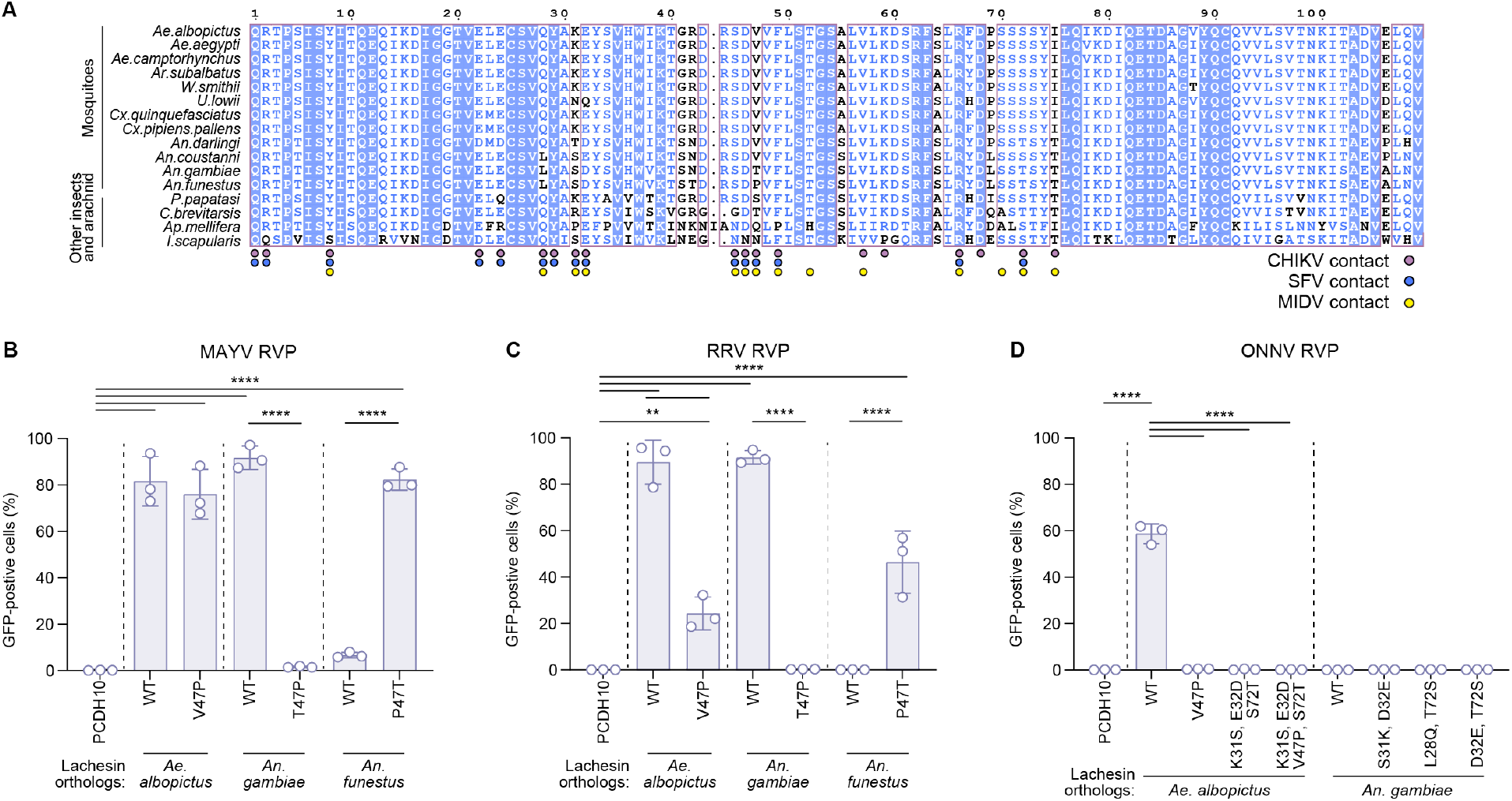
Sequence alignment of Domain 1 (D1) of Lachesin orthologs of mosquitoes and non-mosquito insects, and Lachesin determinants of MAYV, RRV, ONNV binding. **(A)** Sequence alignment of Lachesin D1, generated using ESPript3.0 (*69*). The blue background denotes residues that are completely conserved in all sequences. Boxed residues indicate regions where a single majority residue or multiple chemically similar residues are found, and these residues are shown in blue font. All other residues are shown in black font. CHIKV, SFV, and MIDV contacts are indicated. **(B)** Infection of K562 cells expressing wild-type or mutant *Ae. albopictus, An. gambiae, An. funestus* Lachesin orthologs by GFP-expressing MAYV LET-1430 RVPs at MOI of 1 (measured on K562 cells expressing *Ae. albopictus* Lachesin). Infection was quantified using flow cytometry. **(C)** Infection of K562 cells expressing wild-type or mutant *Ae. albopictus, An. gambiae, An. funestus* Lachesin orthologs by GFP-expressing RRV T48 RVPs at MOI of 1 (measured on K562 cells expressing *Ae. albopictus* Lachesin). Infection was quantified using flow cytometry. **(D)** Infection of K562 cells expressing wild-type or mutant *Ae. albopictus* or *An. gambiae* Lachesin orthologs by GFP-expressing ONNV SG650 RVPs at MOI of 1 (measured on K562 cells expressing *Ae. albopictus* Lachesin). Infection was quantified using flow cytometry. Data are mean ± s.d. from three experiments performed in duplicates (*n* = 3) (**B, C, D**). One-way ANOVA with Šídák’s multiple comparisons test, \*\*\*\**P* < 0.0001 (**B, C, D**), \*\**P* = 0.006 (**C**).

**Table S1.** Cryo-EM data collection and validation statistics.

|  | **CHIKV VLP:**<br>**Lachesin-Fc** | **CHIKV VLP:**<br>**Human MXRA8-Fc** | **SFV VLP:**<br>**Lachesin-Fc** | **MIDV VLP**<br>**Lachesin-Fc** |
| --- | --- | --- | --- | --- |
| EMDB | EMD-77521 | EMD-77522 | EMD-77523 | EMD-77524 |
| PDB ID | 36GF | 36GG | 36GH | 36GI |
| **Data collection** |  |  |  |  |
| Voltage (kV) | 300 | 300 | 300 | 300 |
| Magnification | 130,000 | 130,000 | 130,000 | 130,000 |
| Exposure (e/Å<sup>2</sup>) | 48.7 | 52.6 | 53.4 | 50.6 |
| Defocus range (μm) | -0.8 to -1.8 | -0.8 to -1.8 | -0.8 to -1.8 | -0.8 to -1.8 |
| Pixel size (Å/pixel) | 0.94 | 0.94 | 0.94 | 0.94 |
| **Data processing** |  |  |  |  |
| Final particles | 5,941 | 15,718 | 21,149 | 54,943 |
| VLP resolution (Å) | 6.4 | 4.4 | 4.9 | 4.2 |
| FSC threshold | 0.143 | 0.143 | 0.143 | 0.143 |
| **Block-based re-**<br>**construction** |  |  |  |  |
| Final blocks | 177,416 | 474,538 | 368,721 | 1,658,614 |
| ASU resolution (Å) | 3.4 | 3.0 | 3.2 | 2.9 |
| FSC threshold | 0.143 | 0.143 | 0.143 | 0.143 |
| **Model refinement**<br>**and validation** |  |  |  |  |
| Initial model used | 6NK5/AlphaFold 3 | 8UA8/AlphaFold 3 | 6NK5/6JO8 | AlphaFold 3 |
| R.m.s deviations |  |  |  |  |
| Bond length (Å) | 0.005 | 0.003 | 0.003 | 0.004 |
| Bond angles (°) | 0.887 | 0.751 | 0.660 | 0.640 |
| Validation |  |  |  |  |
| MolProbity score | 1.78 | 1.65 | 1.56 | 1.22 |
| Clash score | 7.34 | 6.09 | 4.20 | 3.12 |
| Ramachandran plot |  |  |  |  |
| Favored (%) | 94 | 96 | 95 | 97 |
| Allowed (%) | 6 | 4 | 5 | 3 |
| Outliers (%) | 0 | 0 | 0 | 0 |

**Table S2.** Buried surface area calculations for alphavirus E2–E1 receptor complexes.

| Complex | BSA (Å<sup>2</sup>) |
| --- | --- |
| CHIKV E2–E1 *Ae. albopictus* Lachesin–Fc | 1,476 |
| SFV E2–E1 *Ae. albopictus* Lachesin–Fc | 1,516 |
| MIDV E2–E1 *Ae. albopictus* Lachesin–Fc | 1,478 |
| CHIKV E2–E1 human MXRA8–Fc | 2,187 |

